# A starvation-triggered AAA+ ATPase halts chromosome replication progression by disassembling the bacterial DNA sliding clamp

**DOI:** 10.1101/2025.01.13.632716

**Authors:** Surbhi, Arnab Kumar Shau, Feby Mariam Chacko, Sunish Kumar Radhakrishnan

## Abstract

Living cells should coordinate vital events such as DNA replication with the availability of nutrients. For example, when cells encounter starvation, to maintain genomic integrity they should harbour robust mechanisms to stop DNA replication. Mechanisms regulating the progression of DNA replication when bacterial cells encounter starvation remain largely unclear. Herein, we identify the role of IncA –a AAA+ ATPase homologous to the prokaryotic RarA and eukaryotic WRNIP1/Mgs1– in inhibiting the progression of chromosome replication in nutrient-starved stationary phase cells of *Caulobacter crescentus*. We show that the starvation-induced alarmone (p)ppGpp ensures the confinement of IncA production to the stationary phase cells. At the mechanistic level, IncA directly interacts with the β-sliding clamp protein DnaN and disassembles DnaN from the replisome, thereby stalling the progression of DNA replication. Furthermore, we reveal the requirement of IncA’s ATPase activity for disassembling DnaN. Remarkably, we demonstrate that the IncA homolog from *E. coli* is capable of inhibiting DNA replication in *Caulobacter*. We propose that IncA homologs serve a stress-dependent role in inhibiting DNA replication across diverse domains of life.

## Introduction

Cells of prokaryotic and eukaryotic origin alike use various mechanisms to regulate vital cellular processes such as chromosome replication in response to nutrient availability. For example, bacterial cells encounter vast fluctuations in nutrient availability warranting the requirement for a robust mechanism to maintain genomic integrity and proliferation under nutrient-limiting conditions. In bacteria, depletion of the nutrient pool leads to the stringent response which regulates several cell cycle events and various developmental processes including DNA replication. The stringent response activates the RelA/SpoT homolog (Rsh) proteins that initiate the production of guanosine tetra- or penta-phosphate molecule, (p)ppGpp (Bange *et al*, 2021). The production of (p)ppGpp influences transcription and translation causing an upregulation of stress-responsive factors and a downregulation of factors involved in growth and development during nutrient-replete conditions (Hauryliuk *et al*, 2015; Irving *et al*, 2021; Travis & Schumacher, 2022; Voedts *et al*, 2024). One of the prominent processes that is affected by the increase in (p)ppGpp levels is chromosome replication.

The bidirectional replication of the circular chromosome in bacteria is divided into initiation, elongation, and segregation events. The initiation event is triggered by the activity of the AAA+ ATPase DnaA (Gorbatyuk & Marczynski, 2001; Sekimizu *et al*, 1987). Upon initiation, the replisome consisting of the helicase (DnaB), the single-stranded DNA-binding protein (SSB), and the DNA polymerase III holoenzyme is loaded onto the replication fork. The DNA polymerase III holoenzyme comprises the core polymerase enzyme and the β-sliding clamp (Aakre *et al*, 2013; Lewis *et al*, 2016). The bacterial β-sliding clamp protein DnaN is important for tethering DNA polymerase III to the replicating DNA and modulating the activity of the polymerase (Altieri & Kelman, 2018). Removal of DnaN leads to the collapse of the replication machinery, which instantaneously arrests DNA replication (Maffeo *et al*, 2019).

In the dimorphic model *Caulobacter crescentus* (henceforth *Caulobacter*), the chromosome replication, and the segregation of the completely replicated chromosome happens only once per cell division cycle. The cell cycle in *Caulobacter* is tightly regulated and consists of a DNA replication-incompetent swarmer cell (G1), which is licensed to initiate chromosome replication only upon entry into the replicative stalked cell phase (S-phase). The primary purpose of the swarmer cell is to disperse off from a nutrient-deprived niche and colonize a new nutrient-rich environment. The swarmer to stalked cell transition (G1 to S) is proposed to be nutrient-dependent (Barrows & Goley, 2023; Britos *et al*, 2011; Kirkpatrick & Viollier, 2012; Skerker & Laub, 2004). Interestingly, the abundance of (p)ppGpp varies during the cell cycle in *Caulobacter*. (p)ppGpp levels are higher in the swarmer cells and decrease upon entry into the replicative stalked cell phase (Boutte *et al*, 2012; Gonzalez & Collier, 2014).

In the swarmer cell, DNA replication is silenced by the overabundance of the master cell cycle regulator CtrA and low levels of ATP-bound DnaA - the DNA replication initiation-competent form of DnaA (Bastedo & Marczynski, 2009; Gorbatyuk & Marczynski, 2005; Quon *et al*, 1998; Sanselicio *et al*, 2015). Upon transition of the swarmer cell into a stalked cell, CtrA is proteolyzed co-incident with the increase in abundance of ATP-bound DnaA which in turn leads to the initiation of chromosome replication (Felletti *et al*, 2019; Quon *et al*., 1998; Ryan *et al*, 2002; Taylor *et al*, 2011). Interestingly, the abundance of CtrA and DnaA in the swarmer cell is influenced by (p)ppGpp (Lesley & Shapiro, 2008; Sanselicio *et al*., 2015). A similar influence of (p)ppGpp on DnaA levels has been demonstrated in *E. coli* (Chiaramello & Zyskind, 1990). However, an overproduction of DnaA is insufficient to trigger DNA replication at (p)ppGpp-abundant conditions (Leslie *et al*, 2015; Riber & Lobner-Olesen, 2020; Sinha *et al*, 2020). Nevertheless, an expression of (p)ppGpp-blind RNA polymerase suffices to initiate replication in nutrient-starved (p)ppGpp-replete conditions (Riber & Lobner-Olesen, 2020). Taken together, these observations allude to the presence of additional factors that are under the control of (p)ppGpp that influence DNA replication in nutrient-deprived conditions.

Herein, we describe a (p)ppGpp-triggered mechanism of DNA replication elongation control in nutrient-starved stationary phase cells of *Caulobacter*. We identify a hitherto uncharacterized function for IncA, which shares homology with the functionally enigmatic AAA+ ATPases RarA (bacteria), Mgs1 (yeast), and WRNIP1 (human), all belonging to the clamp loader clade of proteins (Carrasco *et al*, 2018; Hishida *et al*, 2002; Page *et al*, 2011; Yoshimura *et al*, 2017). We demonstrate that (p)ppGpp transcriptionally favors the production of *incA* specifically in the stationary phase cells. The protein IncA then directly binds to the β-sliding clamp DnaN. The binding of IncA to DnaN inhibits the assembly of DnaN. Furthermore, overexpression of IncA inhibits replication in exponentially growing wild-type cells of *Caulobacter* indicating that the high abundance of IncA is sufficient to inhibit replication. Finally, we show that the ATPase activity of IncA is required for the delocalization of DnaN but not for the binding of IncA to DnaN. Our results identify a conserved nutrient stress-dependent regulator of chromosome replication in bacteria.

## Results

### An overexpression screen identifies incA

Several AAA+ ATPase domain harboring proteins have been shown to orchestrate essential cellular processes in all domains of life. Nevertheless, many remain functionally uncharacterized. Therefore, we performed an overexpression screen using AAA+ ATPase domain harboring proteins that are uncharacterized *Caulobacter*. Herein, we screened for those proteins that decreased viability upon overexpression in *Caulobacter*. Our overexpression screen led us to the identification of CCNA_01342, henceforth named *incA* (<u>in</u>hibitor of <u>c</u>hromosome replication). Overexpression of *incA* from a vanillate inducible promoter on the high copy vector pBVMCS-4 (pP*_van_*-*incA*) decreased the viability of wild-type (*WT*) *Caulobacter* cells (Fig. 1C). Moreover, a 3 h overproduction of *incA* from pP*_van_*-*incA* made the *WT* cells significantly filamentous (Fig. 1A, B and Supplemental Fig. S1A-C). Together, these results suggested that IncA may be involved in regulating a crucial process during the growth of *Caulobacter* cells, and overproduction of IncA could interfere with this process leading to problems in cell cycle and proliferation. From the genome-wide transcriptome data, during the *Caulobacter* cell cycle, it was evident that *incA* transcripts are specifically abundant in the swarmer cells (Schrader *et al*, 2016). This prompted us to investigate the regulatory mechanism that controlled the abundance of IncA in *Caulobacter*.

**Figure 1.**
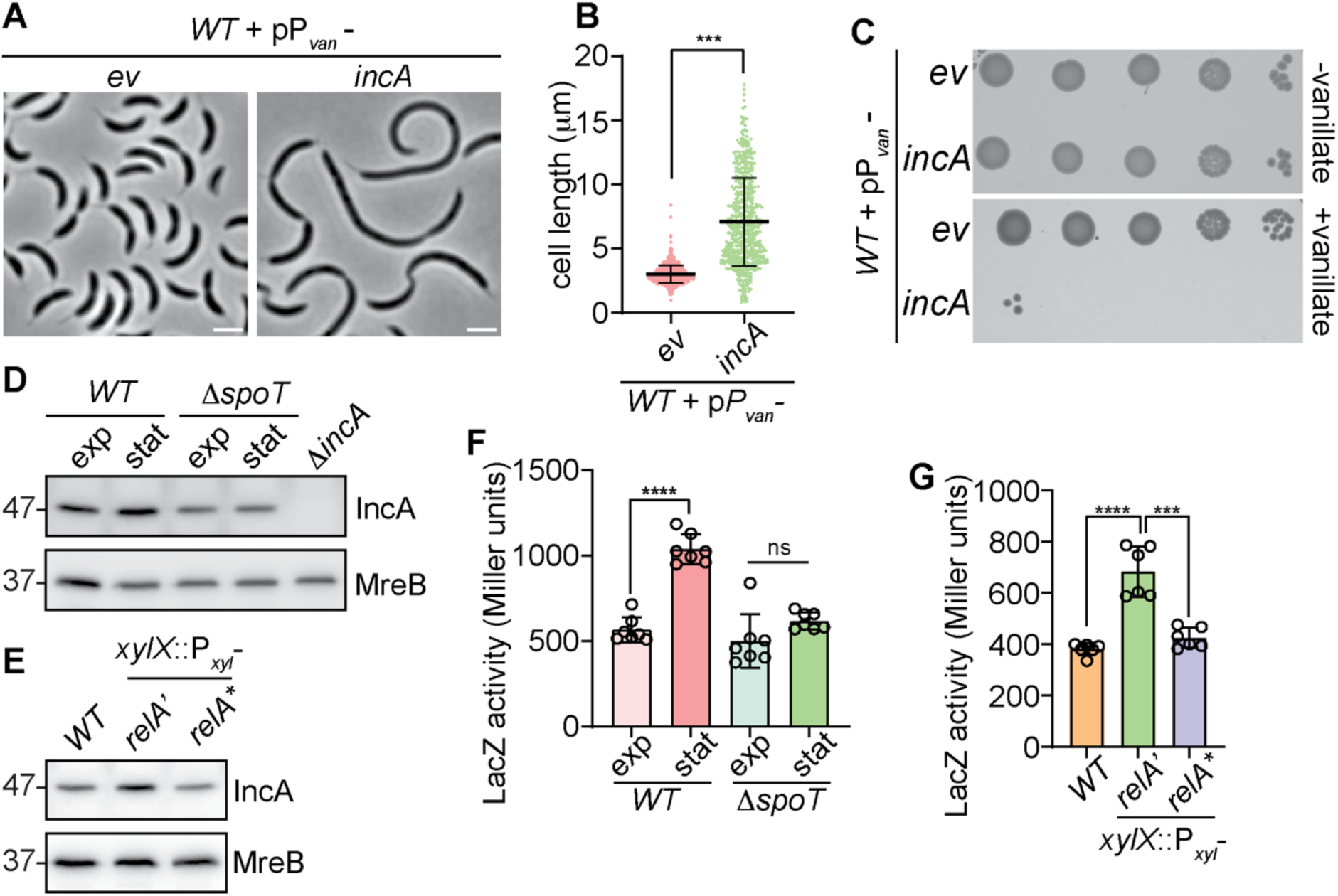
IncA expression is (p)ppGpp-dependent and overexpression reduces cell viability. **(A)** Phase contrast micrographs of wild-type (*WT*) *Caulobacter* cells harbouring the high copy vector (pBVMCS-4) or ectopically expressing *incA* from the vanillate-inducible promoter (P*_van_*) on pBVMCS-4. Cells were induced with 0.5mM vanillate for 5 hours. **(B)** Cell size distribution of cells from (A). Mean cell size of at least 150 cells per biological replicate were used for statistical analyses. **(C)** Growth of cells shown in (A). Ten-fold dilutions of the indicated strains were spotted on to the growth medium with or without the inducer vanillate (0.5 mM). Immunoblots showing the protein levels of IncA and MreB (loading control) in **(D)** *WT* and *spoT* null mutant (Δ*spoT*) during exponential (exp) and stationary (sta) phase and **(E)** exponential phase cells expressing RelA’ or RelA*. Both RelA’ and RelA* were expressed from the xylose-inducible promoter (P*_xyl_*) at the chromosomal *xylX* locus (*xylX*::P*_xyl_*). β-galactosidase (lacZ) activity of the *incA* promoter (P*_incA_*) fused to the lacZ reporter (P*_incA_-lacZ*) in **(F)** *WT* and *ΔspoT* cells at exponential and stationary phase and **(G)** *WT* cells expressing *relA*’ and *relA*\*. The error bars in B, F and G represent the mean ± SD from at least three independent biological replicates. Statistical analyses were done using unpaired two-tailed t test; ****p < 0.0001, ***p < 0.001; ns, not significant. Scale bar: 2μm.

### The stationary phase-specific production of IncA is (p)ppGpp dependent

Interestingly, a study to understand the transcriptional regulatory pattern during stringent response in *Caulobacter* had shown that *incA* transcript levels were up to fourfold less abundant in a *spoT* null (Δ*spoT*) mutant (Boutte & Crosson, 2011). The RelA/SpoT homologs (Rsh) are involved in the production of (p)ppGpp during nutrient stress or at the stationary phase (Bange *et al*., 2021). *Caulobacter* harbors only a single Rsh, *spoT* (Boutte & Crosson, 2011). The Δ*spoT* cells of *Caulobacter* are incapable of making (p)ppGpp at the stationary phase (Ronneau & Hallez, 2019). Therefore, we wondered if the production of *incA* happens at the stationary phase when nutrients are scarce and (p)ppGpp levels are high. To test this, we used polyclonal antibodies generated against IncA to compare the abundance of IncA in exponential and stationary phase cells. Immunoblot analyses indicated that IncA protein levels are indeed more abundant in the stationary phase cells (Fig. 1D). There was almost a two-fold higher amount of IncA at the stationary phase than in the exponential phase cells (Supplemental Fig. S1D). During the cell cycle in *Caulobacter*, (p)ppGpp levels are also found to be abundant in swarmer cells (Boutte *et al*., 2012; Lesley & Shapiro, 2008; Stott *et al*, 2015). This, together with the observation that *incA* transcript levels were abundant in the swarmer cells (Schrader *et al*., 2016), prompted us to wonder if the (p)ppGpp-dependent regulation of *incA* production happens at the level of transcription. To test this, we used a β-galactosidase (*lacZ)*-based reporter fusion wherein the promoter of *incA* was fused to the promoter-less *lacZ* reporter gene (P*_incA_*-*lacZ*). We tested the activity of P*_incA_*-*lacZ* in *WT* cells at exponential and stationary phase conditions. The P*_incA_*-*lacZ* activity was found to be significantly upregulated in the stationary phase cells (Fig. 1F). Moreover, the stationary phase-specific upregulation of the P*_incA_*-*lacZ* activity and the IncA protein abundance were absent in the Δ*spoT* cells (Fig. 1D, F and Supplemental Fig. S1D). Next, we wondered if an artificial increase in (p)ppGpp levels could increase the production of *incA* in *WT* cells even during the nutrient-replete exponential phase conditions. To test this, we used a mutant form of *E. coli* RelA (RelA’) that lacks the C-terminal regulatory domain of RelA (Gropp *et al*, 2001). The absence of the regulatory domain makes RelA’ constitutively active, inducing the production of (p)ppGpp even at normal conditions (Bange *et al*., 2021; Gonzalez & Collier, 2014; Gropp *et al*., 2001). Furthermore, expression of RelA’ in *WT Caulobacter* from the xylose inducible promoter on the chromosome (*xylX::*P*_xyl_*-*relA’*) has been shown to increase the abundance of (p)ppGpp (Gonzalez & Collier, 2014). Immunoblot and P*_incA_*-*lacZ* analyses indicated that the IncA protein levels and the P*_incA_*-*lacZ* activity were indeed increased in cells producing *relA’* from *xylX*::P*_xyl_*-*relA’* (Fig.1E, G and Supplemental Fig. S1E). It has been demonstrated that RelA’ becomes catalytically inactive if the E335 residue at its enzymatically active center is mutated (E335Q) (Gonzalez & Collier, 2014; Harinarayanan *et al*, 2008). There was no difference in the IncA levels or P*_incA_*-*lacZ* activity when *relA’* harbouring the E335Q mutation (*relA\**) was expressed from the *xylX* locus (*xylX::*P*_xyl_*-*relA\**) (Fig.1E, G and Supplemental Fig. S1E) suggesting that it is the (p)ppGpp production by RelA’ that influenced the synthesis of *incA*.

Taken together, our results indicated that IncA production is significantly increased in the stationary phase cells, and that the stationary phase-specific increase of *incA* production was through the SpoT-dependent production of (p)ppGpp.

### IncA interacts with the DNA sliding clamp

To understand the mechanism by which IncA overproduction induces cell filamentation and decreases the viability of *WT Caulobacter* cells, we decided to identify the interacting partners of IncA. Towards this, we carried out co-purification experiments using a C-terminal tandem affinity purification (TAP)-tagged IncA (IncA-TAP) expressed from the vanillate-inducible promoter on the pBVMCS-4 (pP*_van_*-*incA*-*TAP*) in Δ*incA* cells. The co-purified samples were then analyzed using mass spectrometry. The relative abundance of co-purified proteins was quantified against affinity-purified samples from the Δ*incA* cells harboring the empty TAP vector (see methods). From our analyses we found that the replisome component proteins such as the β-sliding clamp DnaN, the nucleoid-associated protein HU, and the single-stranded DNA binding protein SSB co-purified with notable abundance in the IncA-TAP samples (Fig. 2A, Supplemental Dataset 1). Among them, DnaN co-purified with at least two-fold abundance (Fig. 2A, Supplemental Dataset 1).

**Figure 2.**
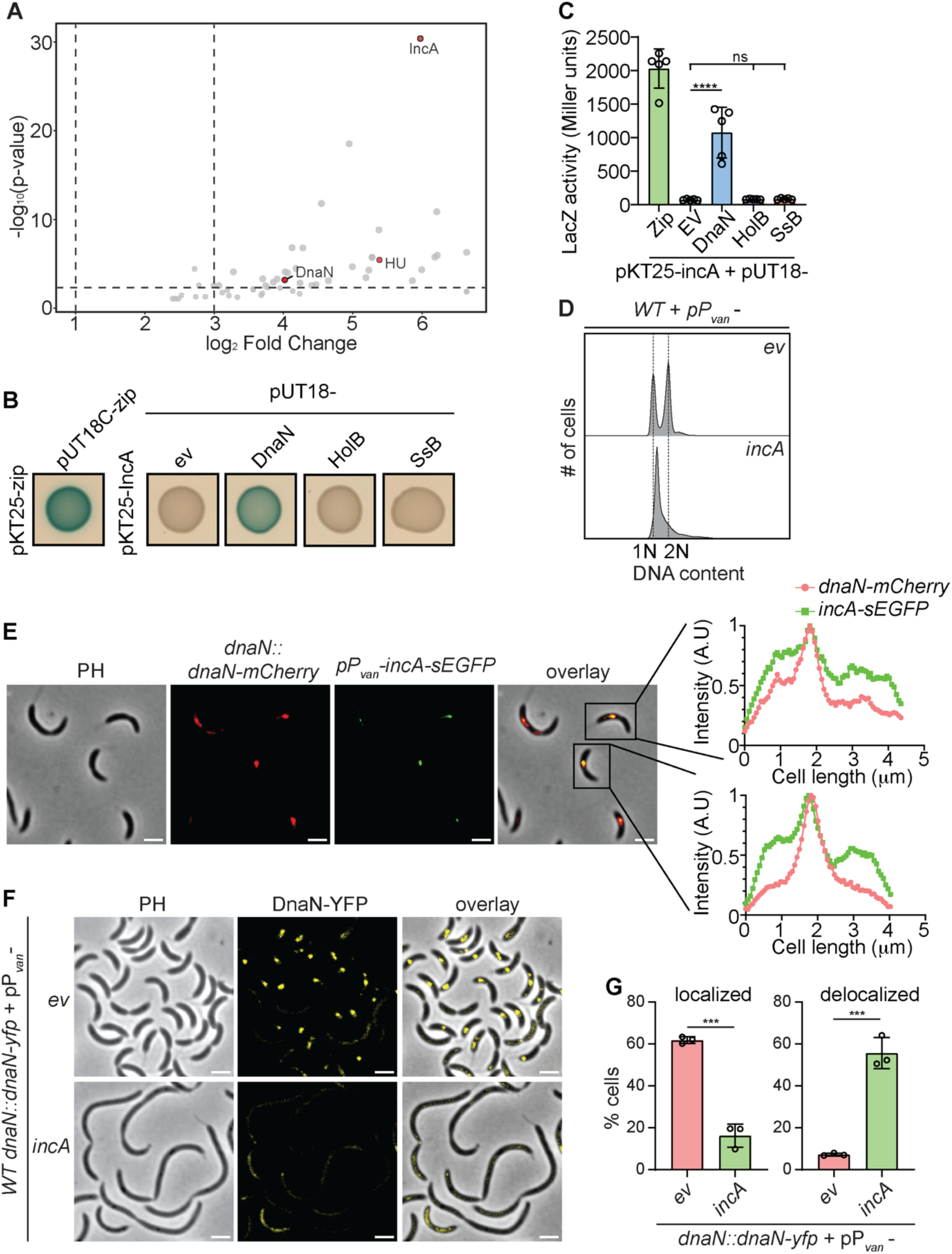
IncA interacts with and delocalizes the DNA sliding clamp DnaN. **(A)** Quantitative proteomics of tandem affinity purified (TAP) samples from *incA*-null mutants expressing the TAP-tag alone or IncA-TAP shown as volcano plots denoting the abundant proteins in the IncA-TAP samples. The threshold was set for log_2_ (fold change) and -log10 (p-value) (see Supplemental Dataset 1). **(B)** Bacterial two-hybrid (BACTH) analysis between IncA fused to T25 fragment of adenylate cyclase at N-terminus (T25-IncA) and replisome components DnaN or HolB or SSB fused to T18 fragment of adenylate cyclase at C-terminus: DnaN-T18; HolB-T18; SSB-T18, respectively. The *E. coli BTH101* cells containing the pair-wise combination of the plasmid constructs were used for the BACTH assay. Plasmids containing leucine zipper motifs of GCN4(pKT24-zip and pUT18C-zip) were used as a positive control. The interaction is denoted by the appearance of blue colour on plates containing X-gal. **(C)** Quantification of interactions in (B) represented as β-galactosidase (LacZ) activity. **(D)** Flow cytometry profiles showing DNA content in wild-type (*WT*) cells harbouring the empty vector pBVMCS-4 (*ev*) or overexpressing *incA* from P_van_ on pBVMCS-4 (pP*_van_-incA*). Cells were treated with 0.5mM vanillate for three hours to induce the expression of *incA*. **(E)** Micrographs denoting the co-localization of DnaN-mCherry and IncA-seGFP. Fluorescence profiles for DnaN-mCherry and IncA-seGFP are plotted for two representative cells. DnaN-mCherry was expressed from the native *dnaN* locus (*dnaN::dnaN-mCherry*) and IncA-seGFP was expressed from P*_van_* on PBVMCS-4 (pP*_van_*-*incA*-*seGFP*). **(F)** Phase contrast and fluorescence micrographs denoting the localization of DnaN-YFP in *WT* cells harbouring the vector pBVMCS-4 (*ev*) or overexpressing *incA* from the vanillate-inducible promoter on pBVMCS-4 (pP*_van_-incA*). DnaN-YFP was expressed from the native *dnaN* locus (*dnaN*::*dnaN*-*yfp*). **(G)** Quantification of localized and delocalized DnaN-YFP in cells from (F) using at least one hundred cells from each biological replicate. The error bars in (C) and (G) represent mean ± SD from at least three independent biological replicates. Statistical analyses were done using a two-way ANOVA with Holm-Sidak’s multiple comparisons test in (C) and unpaired two-tailed t test in (G); ****p < 0.0001, ***p < 0.001; ns, not significant. Scale bar: 2μm.

Furthermore, to confirm if IncA interacts directly with any of the replisome components, we resorted to bacterial-two-hybrid assays using the bacterial adenylate cyclase-based two-hybrid (BACTH) assay system in *E. coli* (Karimova *et al*, 1998). The BACTH system relies on the assisted functional reconstitution of the two fragments, T18 and T25, of the adenylate cyclase from *Bordetella pertussis* to trigger cAMP-dependent LacZ expression (Karimova *et al*., 1998). The BACTH assay indicated that IncA directly interacts with DnaN (Fig. 2B, C) and not with SSB or the DNA polymerase III subunit HolB (Fig. 2B, C). Furthermore, localization experiments using cells producing DnaN-mCherry from the native *dnaN* locus (*dnaN*::*dnaN-mCherry*) and IncA-seGFP from the vanillate-inducible promoter on pBVMCS-4 (pP*_van_*-*incA-seGFP*) indicated that IncA-seGFP co-localized with DnaN-mCherry in 46% of cells with proper DnaN-mCherry foci (Fig. 2E and Supplemental Fig. S2F). The above experiments established the interaction of IncA with DnaN and suggested that the IncA overexpression phenotype may be because of the effect of IncA on DnaN.

### IncA inhibits chromosome replication through DnaN

The β-sliding clamp DnaN is an essential component of the replisome and is required for tethering and increasing the processivity of the DNA polymerase III complex while sliding along the DNA to ensure efficient DNA synthesis (Johnson & O’Donnell, 2005; Yao & O’Donnell, 2021). Inhibition of DnaN leads to replication collapse in actively growing cells (Maffeo *et al*., 2019). From the BACTH experiments, it was evident that IncA directly binds to DnaN (Fig. 2B). Therefore, we wondered if the toxicity in the IncA overexpressing cells is due to the inhibition of DnaN activity by IncA leading to a DNA replication arrest. To test if DNA replication is inhibited in cells overproducing IncA, we resorted to flow cytometry analyses to monitor the replication status. Towards this, we used cells treated with rifampicin. The antibiotic rifampicin is known to specifically inhibit replication initiation while allowing the completion of replication elongation and segregation of replicated DNA. If DNA replication elongation is inhibited by IncA, then proper completion of replication and the segregation of the replicated DNA will be hindered in IncA overexpressing cells. From the flow cytometry analyses properly replicated 1N and 2N chromosome-harboring cells were observed in the rifampicin-treated *WT* harboring the empty vector (Fig. 2D). Strikingly, no such proper 1N and 2N chromosome-harboring cells were observed from the flow cytometry analyses of rifampicin-treated *WT* cells overexpressing *incA* from pP*_van_*-*incA*. This result suggested that DNA replication and segregation were not completed when IncA was overproduced (Fig. 2D). To further test if the chromosome replication has been initiated in the IncA overexpressing cells, we decided to monitor the localization of GFP-ParB. The GFP-tagged version of the chromosome partitioning protein ParB (GFP-ParB) is known to bind to the origin-proximal region and serves as a proxy to monitor replication initiation (Narayanan *et al*, 2018). Due to the duplication of the origin-proximal region upon replication initiation, cells in which chromosome replication has been initiated will display two GFP-ParB foci. We overexpressed IncA in *WT* cells expressing GFP-ParB from the native *parB* locus (*parB::gfp-parB*). Two GFP-ParB foci were observed in 64% of cells overexpressing IncA (Supplemental Fig. S2A) indicating that chromosome replication has been initiated in the IncA overexpressing cells. Inhibition of chromosome replication progression induces SOS response (Maslowska *et al*, 2019). Therefore, we wondered if SOS response is induced in IncA overexpressing cells. To test this, we decided to monitor the activation of the promoter of *sidA*, the SOS-specific inhibitor of cell division in *Caulobacter*, which is known to be activated only upon the induction of SOS response (Modell *et al*, 2011). We overexpressed IncA in *WT* cells expressing YFP from the *sidA* promoter (P*_sidA_*) at the chromosomal xylose (*xyl*) locus in *Caulobacter* (*xylX::*P*_sidA_-yfp*). The production of YFP from P*_sidA_* was increased in cells overexpressing IncA indicating the induction of SOS response in these cells (Supplemental Fig. S2C). Together, these results suggested IncA directly interacts with DnaN and that the overabundance of IncA inhibited progression of DNA replication.

Next, we wondered about influence of IncA on DnaN. Towards this, we overexpressed IncA in cells harboring a functional DnaN-YFP produced from the native *dnaN* locus (*dnaN::dnaN-yfp*). Fluorescence micrographs indicated the presence of intact DnaN-YFP foci in cells harboring the empty vector (Fig. 2F, G). Strikingly, upon overexpression of IncA, DnaN-YFP was delocalized and did not form prominent foci (Fig. 2F, G, Supplemental Movie 1 and 2). Overexpression of IncA led to the delocalization of DnaN-YFP in about 60% of cells (Fig. 2G). Immunoblot analyses indicated that IncA overexpression did not affect the DnaN-YFP protein levels (Supplemental Fig. S2D). Furthermore, immunoblot analyses of co-purified samples from TAP-tagged IncA in DnaN-YFP expressing cells confirmed the *in vivo* interaction of IncA with DnaN-YFP (Supplemental Fig. S2E). Finally, overexpression of IncA also dislodged the delta prime subunit of the DNA polymerase III (HolB) further confirming that the delocalization of DnaN by IncA disrupted the replisome (Supplemental Fig. S2B).

Expression analyses of *incA* indicated that IncA is profoundly produced in the stationary phase cells (Fig. 1D, F). This prompted us to wonder if the role of IncA is to inhibit replication in the stationary phase cells by inhibiting DnaN. To understand this, we localized and quantified DnaN-YFP foci in exponential and stationary phase cells of the *WT* and the *incA* null mutant (Δ*incA*). Proper DnaN-YFP foci were comparable in *WT* and Δ*incA* cells at the exponential phase (Fig. 3A, C). However, at the stationary phase, DnaN-YFP was prominently delocalized in ∼60% of *WT* cells while a majority of Δ*incA* cells had intact DnaN-YFP foci (Fig. 3A, D). To test if this effect is specifically due to the absence of IncA, we ectopically expressed *incA* from its native promoter on a low copy vector (pP*_incA_*-*incA*). Ectopic expression of *incA* in Δ*incA* cells destabilized DnaN-YFP at the stationary phase (Fig. 3B, E) confirming that it is the absence of IncA that allowed intact DnaN-YFP foci in stationary phase cells of Δ*incA*. Deletion of *incA* did not affect the DnaN-YFP protein levels (Supplemental Fig. S3B, C).

**Figure 3.**
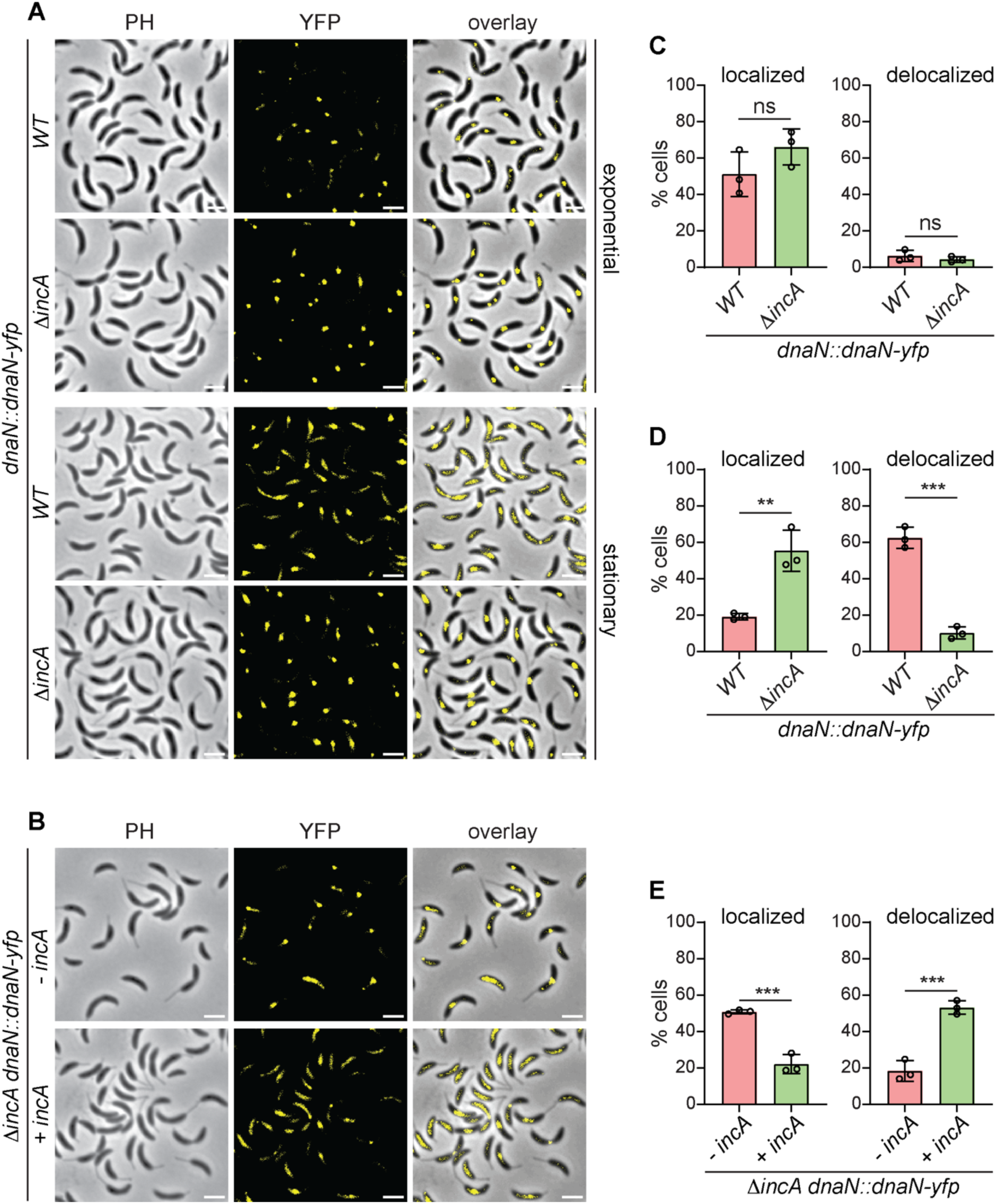
IncA delocalizes DnaN in stationary phase cells. **(A)** Phase contrast and fluorescence micrographs of (A) stationary and exponential phase *WT* and *ΔincA* cells expressing DnaN-YFP from the native *dnaN* locus (*dnaN*::*dnaN*-*yfp*) and **(B)** stationary phase *ΔincA dnaN*::*dnaN*-*yfp* cells expressing *incA* from the *incA* promoter on the low copy vector *plac290*. Exponential phase cells were collected at OD_600_ of 0.6 and stationary phase cells were collected six hours after the exponential phase. (C-E) Quantification (%) of cells from (A) and (B) having localized or delocalized DnaN-YFP. The error bars represent mean ± SD of at least one hundred cells from each of three independent biological replicates. Statistical analyses were done using unpaired two-tailed t test; ***p < 0.001, **p < 0.01; ns, not significant. Scale bar: 2μm.

The production of IncA is under the control of SpoT through (p)ppGpp (Fig. 1D-G). Therefore, we speculated that Δ*spoT* cells should have more intact DnaN-YFP foci due to their inability to produce IncA at the stationary phase. Our analyses indeed indicated the presence of intact DnaN-YFP signals at the stationary phase in Δ*spoT* cells (Supplemental Fig. S3A). Nevertheless, the levels of DnaN-YFP were found to be reduced in Δ*spoT* cells at the stationary phase (Supplemental Fig. S3B), which could be attributed to the pleiotropic effect that cells may encounter in the absence of (p)ppGpp (Boutte & Crosson, 2011; Boutte *et al*., 2012).

Taken together, our analyses bolstered the idea that IncA directly interacts with DnaN. This interaction delocalizes DnaN specifically in the stationary phase cells that have an increased abundance of IncA.

### The ATPase activity of IncA is required to delocalize DnaN

Moving forward, we set out to understand the role of the ATPase domain in IncA towards its activity on DnaN. The aspartate residue in the Walker B motif is a part of the active site in the ATPase domain. The conserved aspartate coordinates with the Mg^2+^ to neutralize the negative charge of phosphate present in ATP. Mutations in the conserved aspartate residue eliminate the ATPase activity while maintaining ATP binding (Puchades *et al*, 2020). Therefore, we mutated the corresponding aspartate residue at the 115^th^ (D115) position on IncA to alanine (IncA^D115A^) (Supplemental Fig. S4A). Overexpression of *incA*^D115A^ from the vanillate inducible promoter on pBVMCS-4 (pP*_van_*-*incA*^D115A^) in Δ*incA* cells induced mild filamentation (Fig. 4A, C, Supplemental Fig. S4B) but did not decrease the viability (Fig. 4D). Moreover, overexpression of IncA^D115A^ did not delocalize DnaN-YFP (Fig. 4A, B, Supplemental Fig. S4C) or inhibit the chromosome replication (Fig. 4E). It is possible that the ATPase activity is required to dislodge DnaN and does not have a role in determining its interaction with DnaN, which we tested using BACTH. Interestingly, our BACTH analyses indicated that IncA^D115A^ could still interact with DnaN (Fig. 4F, G). Together, these experiments suggested that the ATPase mutant of IncA could interact with DnaN but cannot delocalize DnaN-YFP or inhibit replication, indicating that the ATPase activity is required for delocalizing DnaN from the replisome.

**Figure 4.**
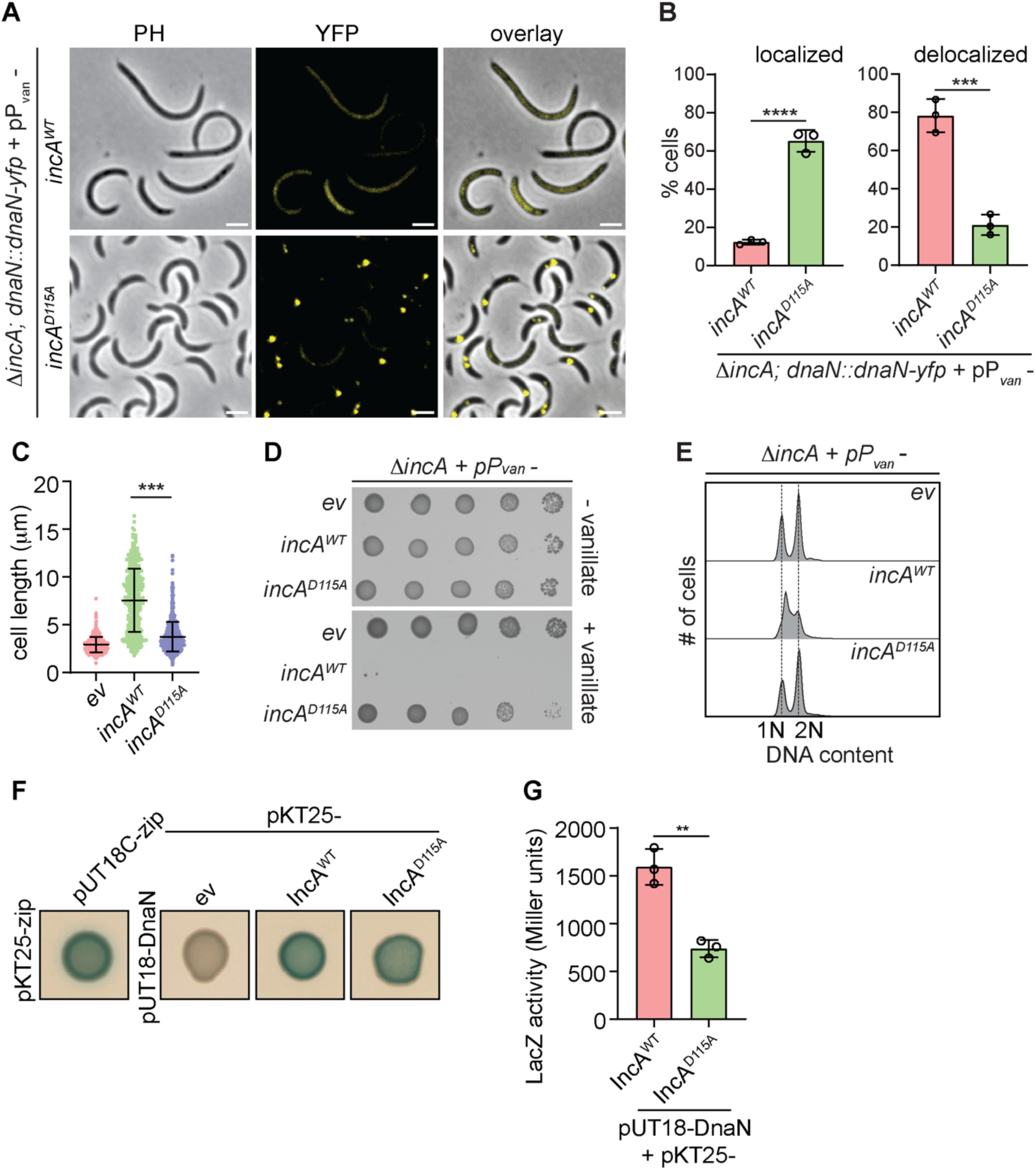
The ATPase activity of IncA is required to inhibit DNA replication. **(A)** Phase contrast and fluorescence micrographs representing DnaN-YFP in *ΔincA* cells overexpressing wild-type IncA (*incA^WT^*) or Walker-B mutant of IncA (*incA^D115A^*) from a vanillate-induble promoter on pBVMCS-4 (pP*_van_*). **(B)** Quantification of cells from (A) denoting the percentage of cells displaying localized and delocalized DnaN-YFP. At least one hundred cells from each of three independent biological replicates were used for quantification. **(C)** Cell size distribution, **(D)** growth and **(E)** Flow cytometry profiles of *ΔincA* cells harbouring the empty vector pBVMCS-4 (*ev*) or overexpressing *incA^WT^* or *incA^D115A^* from P*_van_* on pBVMCS-4 (pP*_van_*). For growth assay, ten-fold dilutions of the indicated strains were spotted onto a growth medium with or without the inducer vanillate (0.5 mM). **(F)** Bacterial two-hybrid (BACTH) analysis between DnaN fused to the T18 fragment of adenylate cyclase at the C-terminus (DnaN-T18) and IncA^WT^ or IncA^D115A^ fused to T25 fragment of adenylate cyclase at the N-terminus: T25-IncA^WT^, T25-IncA^D115A^, respectively. The *E. coli BTH101* cells containing the pair-wise combination of plasmid constructs were used for the BACTH assay. Plasmids containing leucine zipper motifs of GCN4(pKT24-zip and pUT18C-zip) were used as a positive control. The interaction is denoted by the appearance of blue colour on plates containing X-gal. **(G)** Quantification of interactions in (F) represented as β-galactosidase (LacZ) activity. The error bars represent mean ± SD from at least three independent biological replicates. Statistical analyses in (B), (C) and (G) were done using unpaired two-tailed t test; ****p < 0.0001, ***p < 0.001, **p < 0.01; ns, not significant. Scale bar: 2μm.

### The E. coli homolog of IncA inhibits DNA replication in Caulobacter

IncA from *Caulobacter* shares 48% identity with RarA from *E. coli*. Furthermore, AlphaFold and ChimeraX-based structural alignment of monomeric IncA and RarA indicated that IncA shares a significant structural homology with RarA (Fig. 5F). Therefore, we wondered if RarA from *E*. *coli* has a similar function as IncA in *Caulobacter*. To understand this, we overexpressed *E. coli* RarA in Δ*incA* cells of *Caulobacter* from the vanillate inducible promoter on pBVMCS-4 (pP*_van_*-*rarA*). Interestingly, overexpression of RarA induced severe filamentation (Fig.5 A, C) and viability defects (Fig. 5D) in *Caulobacter*. Furthermore, the overexpression of RarA partially delocalized DnaN-YFP (Fig. 5A, B, Supplemental Fig. S5C). Flow cytometry analyses of rifampicin-treated RarA overexpressing *Caulobacter* cells indicated the inhibition of chromosome replication in these cells (Fig. 5E). The observation of DNA replication inhibition was further strengthened by the induction of the SOS response in RarA overexpressing *Caulobacter* cells (Supplemental Fig. S2C). Taken together, our results suggested that the IncA homolog from *E. coli*, RarA, harbors an IncA-like function in *Caulobacter*.

**Figure 5.**
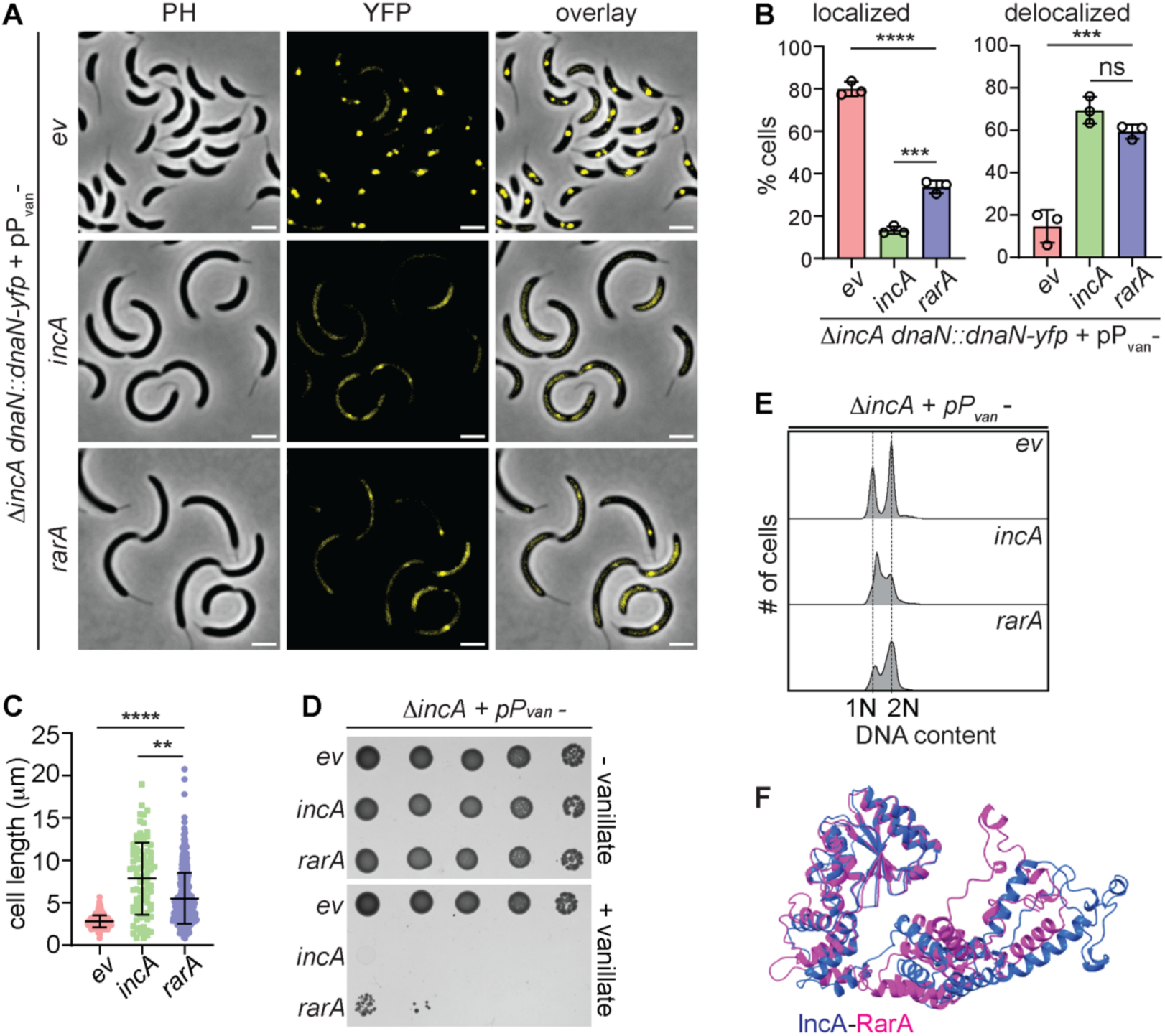
RarA inhibits DNA replication in *Caulobacter*. **(A)** Phase contrast and fluorescence micrographs representing DnaN-YFP in *ΔincA* cells harbouring the vector pBVMCS-4 (*ev*) or overexpressing *Caulobacter incA* or *E. coli rarA* from the vanillate-inducible promoter on pBVMCS-4 (pP*_van_*). **(B)** Quantification of cells from (A) denoting the percentage of cells displaying localized and delocalized DnaN-YFP. At least one hundred cells from each biological replicate was used for quantification. **(C)** Cell size distribution, **(D)** growth and **(E)** flow cytometry profiles of *ΔincA* cells harbouring the empty vector pBVMCS-4 (*ev*) or overexpressing *incA* or *rarA.* For growth assay, ten-fold dilutions of the indicated strains were spotted onto a growth medium with or without the inducer vanillate (0.5 mM). **(F)** Structural comparison of the IncA monomer (Purple) and RarA monomer (Pink). The monomeric structures were generated using Alphafold and docked using UCSF ChimeraX. The error bars in (B) and (C) represent mean ± S.D from at least three independent biological replicates. Statistical analyses were done using unpaired two-tailed t test; ****p < 0.0001, ***p < 0.001; ns, not significant. Scale bar: 2μm.

## Discussion

Free-living cells often encounter nutrient-dependent stress that significantly impacts their survival and adaptability. When essential nutrients become scarce, cells modulate their metabolic processes, often entering a dormant state until nutrients become replete. In bacteria, the starvation response is escalated with the production of the alarmone (p)ppGpp, which affects several cell cycle and developmental processes including chromosome replication (Bange *et al*., 2021; Hauryliuk *et al*., 2015; Ronneau & Hallez, 2019). Nutrient-poor conditions could cripple nucleotide availability which could in turn cause DNA lesions. Therefore, to prevent damage to the genetic material, it becomes imperative for cells to stop replication at starvation. The mechanisms that bacterial cells utilize to regulate DNA replication at nutrient-starved conditions are not well understood. Herein, we demonstrate the role of IncA, a highly conserved AAA+ ATPase homolog belonging to the clamp loader family, in inhibiting DNA replication at nutrient-starved stationary phase (Fig. 6).

**Figure 6.**
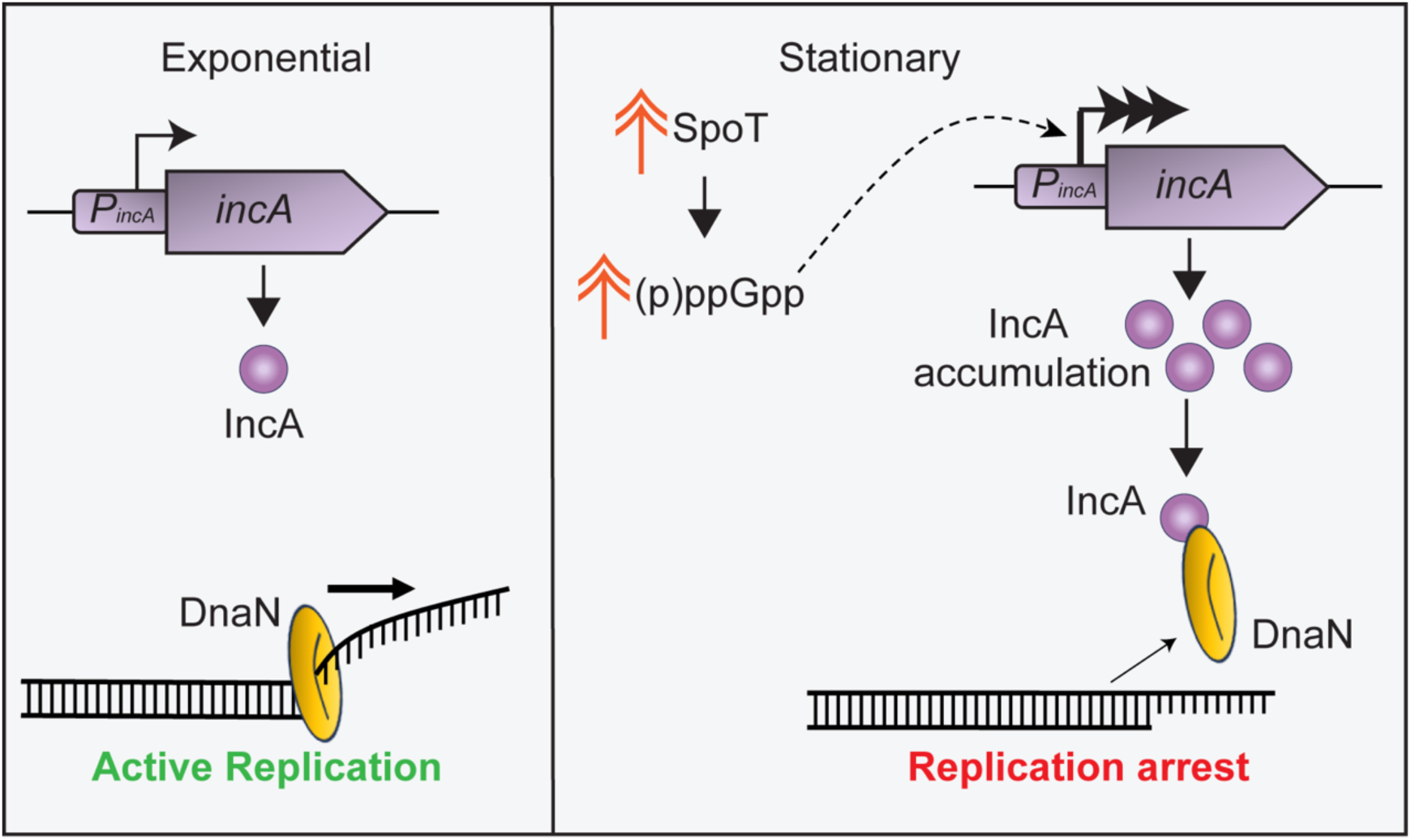
Mechanism of IncA-dependent inhibition of DNA replication. At stationary phase, nutrient starvation enhances the activity of SpoT causing an increase in the abundance of (p)ppGpp. The increased (p)ppGpp levels license the activation of the *incA* promoter (P*_incA_*) leading to an accumulation of the IncA protein, which then directly binds to, and delocalizes, the DNA sliding clamp causing replication arrest. At exponential phase, P*_incA_* remains less active due to the absence of (p)ppGpp.

### The stationary phase-specific transcriptional regulation of incA by (p)ppGpp

The transcription of IncA is tightly linked to the levels of (p)ppGpp whose production is specifically enhanced during starvation. An increase in the (p)ppGpp level suffices to activate the promoter of *incA* leading to IncA production (Fig. 1), suggesting the existence of a direct (p)ppGpp-dependent regulatory module controlling the production of *incA*. This module could work either by directly regulating the RNA polymerase activity or through the regulation of the activity of a transcriptional factor as has been shown in *E. coli* and *Caulobacter* (Bange *et al*., 2021; Boutte & Crosson, 2011; Irving *et al*., 2021; Voedts *et al*., 2024). By coupling *incA* production to (p)ppGpp, *Caulobacter* cells restrict the increased abundance of IncA to nutrient-poor conditions. Thereby, cells ensure that the inhibitory effects of IncA are confined only to starvation-induced stress. Such a type of control could check unwanted replication-induced damage in nutrient-poor conditions wherein carbon and nitrogen sources may become limited leading to a decrease in nucleotide availability (Wang *et al*, 2020).

The IncA-dependent effect on chromosome replication may also be prevalent in the swarmer cells. Swarmer cells of *Caulobacter* are replication-incompetent and stay in a G1-like state until they encounter nutrient-favorable conditions (Barrows & Goley, 2023). The inhibition of replication initiation in the swarmer cells is primarily mediated through an over-abundance of CtrA, which binds to the chromosomal origin to inhibit replication initiation (Quon *et al*., 1998). Interestingly, (p)ppGpp levels are enhanced in the replication incompetent swarmer cells of *Caulobacter* (Boutte *et al*., 2012; Gonzalez & Collier, 2014). Furthermore, from the cell cycle RNA abundance data, it is evident that *incA* levels are enriched in the swarmer cells (Schrader *et al*., 2016). Therefore, it is likely that *Caulobacter* cells might utilize *incA* as an additional control to ensure inhibition of replication in swarmer cells until favorable nutrient conditions are encountered leading to the initiation of S-phase. A direct (p)ppGpp-dependent control of replication elongation has been shown in *Bacillus subtilis*. In *Bacillus*, under nutrient starvation, (p)ppGpp directly binds to the primase (DnaG) resulting in the inhibition of its primer extension activity that leads to stalling of replication elongation (Giramma *et al*, 2021; Wang *et al*, 2007). Nevertheless, similar starvation-dependent replication elongation control has not been reported in Gram-negative bacteria.

### Replisome disassembly by a clamp loader clade protein

The process of DNA replication is carried out by a multiprotein complex, the replisome. The homodimeric β-sliding clamp DnaN plays a crucial role in replication progression by tethering the DNA polymerase complex and sliding along the replicating DNA (Simonsen *et al*, 2024). Removal of DnaN could lead to destabilization of the polymerase complex and collapse of replication (Maffeo *et al*., 2019; Simonsen *et al*., 2024). IncA directly binds to, and delocalizes, DnaN thereby arresting DNA replication progression (Fig. 2). The presence of cells harboring partially replicated chromosomes upon IncA overexpression (Fig. 2E), supports this possibility.

A mutation in the aspartate residue of the Walker B motif, that could affect the ATPase activity of IncA (IncA^D115A^), renders IncA incapable of inhibiting DNA replication (Fig. 4A-D). Nevertheless, the IncA^D115A^ mutant could bind to DnaN (Fig. 4E-F). Therefore, it is conceivable that the ATPase activity in IncA is required to dislodge DnaN from the active replisome after the binding of IncA to DnaN. We speculate that the ATPase activity could be required for IncA to slide along with DnaN or for its disassembly. However, the molecular mechanism that IncA utilizes to delocalize DnaN remains to be resolved.

IncA shares homology with AAA+ ATPases like RarA (Replication associated recombination A) from *E. coli* (Page *et al*., 2011) (Fig. 5F), Mgs1 (Maintenance of genomic stability 1) from *S. cerevisiae* (Hishida *et al*., 2002) (Supplemental Fig. S5A) and WRNIP1 (Werner helicase-interacting protein 1) from humans (Yoshimura *et al*., 2017) (Supplemental Fig. S5B). These AAA+ ATPases belong to the clamp-loader clade of proteins (Ammelburg *et al*, 2006; Barre *et al*, 2001; Erzberger & Berger, 2006; Iyer *et al*, 2004). In *E*. *coli* RarA interacts with SSB and is speculated to be involved in resolving stalled replication forks (Page *et al*., 2011). Nevertheless, it has been suggested that RarA could have a loading or unloading function on ring proteins such as the sliding clamp (Sherratt *et al*, 2004). Interestingly, overexpression of RarA from *E. coli* in *Caulobacter* induces replication arrest and decreased viability (Fig. 5). Furthermore, IncA and RarA overexpression upregulates the SOS response in *Caulobacter*, which is a hallmark of replication collapse (Janion, 2008; Maslowska *et al*., 2019) (Supplemental Fig. S2C). These observations suggest that RarA possesses a DNA replication-inhibitory activity in *Caulobacter*.

It has not escaped our attention that Mgs1 from budding yeast is structurally similar to the replication factor C (RFC), a eukaryotic clamp loader (Hishida *et al*, 2006). Furthermore, during DNA damage stress, the ubiquitination of the eukaryotic sliding clamp PCNA facilitates the binding of Mgs1 to PCNA (Saugar *et al*, 2012). The binding of Mgs1 to the ubiquitinated PCNA disrupts the interaction of PCNA with polymerase δ, leading to the disassembly of the replisome (Branzei *et al*, 2002; Saugar *et al*., 2012). In light of this, we speculate that Mgs1 might harbour an IncA-like effect on PCNA and may have a role in directly disassembling PCNA during stress, which warrants investigation.

## Acknowledgments

Anjana Badrinaraynan, Justine Collier, Vikas Jain, Sean Murray and Manjula Reddy for strains. Microscopy, Flow Cytometry and DST-FIST supported Proteomics Facility (SR/FST/LSII-043/2026) at IISER Pune. Research Fellowships from CSIR to Surbhi (09/0936(11651)/2021-EMR-1) and from IISER Pune to AKS. This work is supported by funds from the DBT-Wellcome Trust India Alliance through a Senior Fellowship to SKR (IA/S/20/2/505202).

## Author contributions

Surbhi and SKR conceptualised and designed the study. Surbhi, AKS and FMC performed experiments and collected data. Surbhi, AKS and SKR analysed data. SKR procured funds. Surbhi and SKR wrote the manuscript.

## Materials and Methods

### Bacterial strains and growth media

*Caulobacter crescentus* NA1000 (Evinger & Agabian, 1977) and derivatives were grown on rich peptone yeast extract (PYE) media containing 0.2% peptone, 0.1% yeast extract, 1 mM MgSO_4_, 0.5 mM CaCl_2_ and incubated at 29 °C, unless specifically mentioned. To induce expression from the P*_van_* promoter, media was supplemented with 0.5 mM vanillate for 3-5 h. The *E. coli* strains were grown in lysogeny broth (LB) media or minimal-A media containing 10.5 g l^-1^ K_2_HPO_4_, 4.5 g l^-1^ KH_2_PO_4_, 1 g l^-1^ (NH_4_)_2_SO_4_, 0.5 g l^-1^ sodium citrate dihydrate, 0.2% Glucose, 1 mM MgSO_4_, 0.0001% thiamine (Miller, 1992) at 37 °C unless specifically mentioned. Antibiotics were used at the following concentrations: kanamycin 20 μg ml^-1^ in solid medium and 5 μg ml^-1^ in liquid (50 μg ml^-1^ for *E. coli*), gentamicin 2.5 μg ml^-1^ in solid medium and 1 μg ml^-1^ in liquid (25 μg ml^-1^ for *E. coli*), tetracycline 1 μg ml^-1^ (10 μg ml^-1^ for *E. coli*), spectinomycin 30 μg ml^-1^ in solid medium and 25 μg ml^-1^ in liquid, ampicillin 100 μg ml^-1^ (for *E.coli*). Plasmids were introduced into *C. crescentus* by electroporation. Strains and plasmids used in this study are listed in Supplemental Tables S1-S3.

### Viability assay

Overnight cultures of NA1000 containing vectors expressing IncA, IncA^D115A^ or RarA from the vanillate inducible promoter (P*_van_*) on the vector pBVMCS-4 (Thanbichler *et al*, 2007) were normalized to OD_600_ of 0.1 before being serially diluted. Aliquots (3 µl) of the dilutions were spotted onto PYE agar containing gentamycin (2.5 μg ml^-1^), with or without 0.5 mM vanillate. Plates were incubated at 30 °C for 2-3 days and imaged using Amersham ImageQuant 800 imager (Cytiva, USA)..

### IncA protein purification and antibody production

To produce antibodies against IncA, the IncA protein with a N-terminal hexahistidine tag (His_6_-IncA) was expressed in *E. coli BL21(DE3)* cells. Cells were then harvested by centrifugation, and the recombinant protein was purified using Ni-NTA (nickel-nitrilotriacetic acid) resin (Qiagen). 0.5 mM IPTG (isopropyl-β-D-thiogalactoside) was used for overexpression of His_6_-IncA.. The purified protein was then used to immunize New Zealand white rabbits (Bioklone Biotech, Chennai, India).

### Microscopy

Phase contrast and fluorescence microscopy were performed using an Olympus IX83 inverted microscope (Evident Scientific, Japan) equipped with a U Plan X Extended Apo 100X (1.45 numerical aperture) objective and an ORCA-Flash4.0 V3 sCMOS camera (Hamamatsu, Japan). Cells were placed on a 1% agarose pad, or 1% agarose pad supplemented with PYE, for imaging. The images were processed and analysed using the Fiji image analysis platform (Schindelin *et al*, 2012).

### Immunoblot analyses

For immunoblot analyses *Caulobacter* cells were pelleted and resuspended in 1X SDS buffer (Tris-Cl (62.5 mM, pH 6.8), SDS (2%), Glycerol (10%), Bromophenol blue (0.004%), β-Mercaptoethanol (2.5%)). The proteins were resolved on Sodium Dodecyl sulfate-polyacrylamide (SDS-PAGE) gel and blotted on to polyvinylidene difluoride (PVDF) Immobilon-P membranes (Merck Millipore). The membranes were then blocked in with TBST solution containing 20 mM Tris-HCl [pH 7.5], 150 mM NaCl, 0.2% Tween-20 and 5% non-fat dry milk followed by incubation with primary antibodies, anti-IncA (1:10,000), anti-MreB (1:30,000) (Figge *et al*, 2004), anti-GFP (1:10,000) (Living Colors JL-8, Clontech Laboratories, CA, USA). The blots were then washed with TBS solution (20 mM Tris-HCl [pH 7.5], 150 mM NaCl), and detected with donkey anti-rabbit or donkey anti-mouse secondary antibodies conjugated to horseradish peroxidase (Jackson ImmunoResearch, USA). The blots were visualized using Amersham ImageQuant 800 imager (Cytiva, USA) after treating with clarity western ECL substrate (Bio-Rad Laboratories, USA).

### Bacterial adenylate cyclase two-hybrid (BACTH) Assay

BACTH system (Euromedex, France) was used to study the protein-protein interactions. The plasmids were constructed and co-transformed in *E. coli* BTH101 Δ*cya* competent cells as per manufacturer’s instructions. This system takes advantage of the catalytic domain of adenylate cyclase (CyaA) from *Bordetella pertussis*, which consists of two inactive fragments, T25 and T18, when separated. The heterodimerization of hybrid-proteins leads to the reassembly of the functional enzyme, enabling cAMP synthesis leading to *lacZ* gene expression. The proteins IncA, IncA^D115A^, DnaN, HolB, SSB, and RarA were cloned into pKT25 or pUT18 plasmids allowing fusion with T25 or T18 fragment of adenylate cyclase. Different combinations of T18 and T25 fusion plasmids were co-transformed in *E. coli BTH101* Δ*cya* cells. Overnight cultures of *BTH101* containing co-transformants were spotted (3 μl) onto LB kanamycin (50 μg ml^-1^), ampicillin (100 μg ml^-1^), IPTG (0.5 mM) and X-Gal (40 μg ml^-1^) plates. The plates were incubated at 30 °C for 24 h and then at 4 °C for few hours, at which point the plates were imaged. The LacZ activity was measured after the same overnight cultures were diluted in fresh LB media containing 0.5 mM IPTG, using the protocol mentioned below for promoter activity assay.

### Flow cytometry

For flow cytometry analyses (Siwach *et al*, 2021), *Caulobacter* cells were grown in the presence of 0.5 mM vanillate for 3 h. Cells were then exposed to 20 μg ml^-1^ of rifampicin for three h to stop initiation of DNA replication. 10 μl of this rifampicin treated cells were then fixed in 1.5 ml of 70% ice-cold ethanol. The samples were then vortexed and stored overnight at -20 °C. 500 μl of these samples were pelleted at 8000 rpm for 5 min. The pellets were washed thrice with 1 ml of FACS-staining buffer (10 mM Tris-HCl (pH-7.2), 1 mM EDTA, 50 mM sodium citrate, 0.01%Triton-X-100) by pelleting at 8000 rpm for 3 min. The washed pellets were resuspended in 1 ml of FACS-staining buffer containing 0.1 mg ml^-1^ RNase A (Roche, Switzerland), followed by incubation at 25 °C for 1 h. Cells were then pelleted at 8000 rpm for 3 min and resuspended in 1 ml FACS-staining buffer containing 0.5 μM SYTOX green nucleic acid stain (Invitrogen, USA) and incubated for 5 min in dark. The stained cells were then analyzed using BD Accuri C6 flow cytometer (BD Biosciences, CA, USA). The flow cytometry data was analysed using the FlowJo software (BD Biosciences).

### Tandem affinity purification

The Tandem affinity purification experiments were carried out as described previously(Narayanan *et al*, 2015). Exponential phase cells (1 l) were harvested by centrifugation at 7000 rpm for 15 min and washed in buffer-1 (50 mM sodium phosphate [pH-7.4], 50 mM NaCl, 1 mM EDTA). Cells were then pelletised and resuspended in buffer-2 (buffer-1 containing 10 mM MgCl_2_, 0.5% n-dodecyl-β-D-maltoside (Sigma-Aldrich), 1x protease inhibitors (Complete EDTA-free, Roche, Switzerland) and lysed by addition of 5000 U of Ready-Lyse Lysozyme Solution (Biosearch Technologies, USA), followed by sonication at 60% amplitude, to reduce the culture viscosity. The sonicated lysate was then centrifuged at 10,500 rpm for 45 min at 4 °C to remove cell debris. The supernatant was then incubated with IgG Sepharose beads (Cytiva, USA) for 2 h at 4 °C with constant mixing. The beads were then washed thrice with IPP150 buffer (10 mM Tris-HCl [pH-8], 150 mM NaCl, 0.1% NP-40) and once with TEV-cleavage buffer (IPP150 buffer containing 0.5 mM EDTA and 1 mM DTT). The washed beads were then incubated overnight at 4 °C with TEV cleavage buffer containing 100 U ml^-1^ TEV protease (Promega, USA), with constant mixing, to release the tagged complex. 3 µM CaCl_2_ was added to the eluate and incubated with Calmodulin-Sepharose beads (Cytiva, USA) for 1 h with constant mixing. The beads were then washed thrice with calmodulin binding buffer (10 mM β-mercaptoethanol, 10 mM Tris-HCl (pH-8), 150 mM NaCl, 1 mM magnesium acetate, 1 mM imidazole, 2 mM CaCl_2_, 0.1% NP-40). The bound proteins from the beads were then eluted using calmodulin elution buffer (10 mM β-mercaptoethanol, 10 mM Tris-HCl (pH-8), 150 mM NaCl, 1 mM magnesium acetate, 1 mM imidazole, 2 mM EGTA, 0.1% NP-40). The eluate was then concentrated using Amicon ultra centrifugal filters, 3 kDa MWCO (Merck-Millipore) and were analysed by immunoblots or mass spectrometry.

### Quantitative mass spectrometry

For quantitative mass-spectrometry (Kamat *et al*, 2015) 5 μg each of the TAP samples was processed for chloroform-methanol precipitation of proteins using 1:1:3 ratio of protein, chloroform, and methanol. The resultant white precipitate, was collected by centrifugation at 20,000 × *g*for 20 mins at 4 °C. The pellet was dried completely using a CentriVap vacuum concentrator (Labconco, USA). The dried pellet was resuspended in 50 μl of 8 M urea in 100 mM tetraethylammonium bicarbonate (TEAB) and sonicated using an ultrasonic cleaner (Spire, India) for 5 min. The degraded proteins were reduced using 5 mM dithiothreitol (DTT), followed by incubation at 60 °C for 30 min. The reaction was cooled down to room temperature before the addition of 15 mM iodoacetamide and incubated in dark for 15 min to alkylate the free sulfhydryl groups of cysteine residues. The mixture was supplemented with an additional 10 mM DTT and the urea was diluted by making up the volume of the reaction to 400 μl using 100 mM TEAB. The reaction was incubated at 37 °C for 16 h with constant shaking after addition of 4 μl of 0.5 mg ml^-1^ trypsin (Promega). After trypsin digestion, the samples were either labelled with 8 μl of 4% HCHO (light formaldehyde) or 8 μl of 4% DCDO (heavy formaldehyde). 8 μl of NaBH_3_CN was added to both reactions and were incubated at room temperature with constant shaking for 1 h, to promote labelling. The reactions were then quenched by the addition of 32 μl of 1% ammonia. Finally, the reactions were stopped by adding 5% formic acid and the supernatant was collected after centrifugation at 20,000 × *g* for 20 min. An equal volume of heavy-labelled and light-labelled peptide samples were mixed and processed for desalting via Empore C18 disks (Merck-Supelco). The samples were then subjected to LC-MS analysis using TripleTOF 6600 System (Sciex, USA). The data was then analysed using the ProteinPilot software (Sciex, USA) and plotted using VolcaNoseR (Goedhart & Luijsterburg, 2020).

## References

Aakre CD, Phung TN, Huang D, Laub MT (2013) A bacterial toxin inhibits DNA replication elongation through a direct interaction with the beta sliding clamp. Mol Cell 52: 617–628

Altieri AS, Kelman Z (2018) DNA Sliding Clamps as Therapeutic Targets. Front Mol Biosci 5: 87

Ammelburg M, Frickey T, Lupas AN (2006) Classification of AAA+ proteins. J Struct Biol 156: 2–11

Bange G, Brodersen DE, Liuzzi A, Steinchen W (2021) Two P or Not Two P: Understanding Regulation by the Bacterial Second Messengers (p)ppGpp. Annu Rev Microbiol 75: 383–406

Barre FX, Soballe B, Michel B, Aroyo M, Robertson M, Sherratt D (2001) Circles: the replication-recombination-chromosome segregation connection. Proc Natl Acad Sci U S A 98: 8189–8195

Barrows JM, Goley ED (2023) Synchronized Swarmers and Sticky Stalks: Caulobacter crescentus as a Model for Bacterial Cell Biology. J Bacteriol 205: e0038422

Bastedo DP, Marczynski GT (2009) CtrA response regulator binding to the Caulobacter chromosome replication origin is required during nutrient and antibiotic stress as well as during cell cycle progression. Mol Microbiol 72: 139–154

Boutte CC, Crosson S (2011) The complex logic of stringent response regulation in Caulobacter crescentus: starvation signalling in an oligotrophic environment. Mol Microbiol 80: 695–714

Boutte CC, Henry JT, Crosson S (2012) ppGpp and polyphosphate modulate cell cycle progression in Caulobacter crescentus. J Bacteriol 194: 28–35

Branzei D, Seki M, Onoda F, Enomoto T (2002) The product of Saccharomyces cerevisiae WHIP/MGS1, a gene related to replication factor C genes, interacts functionally with DNA polymerase delta. Mol Genet Genomics 268: 371–386

Britos L, Abeliuk E, Taverner T, Lipton M, McAdams H, Shapiro L (2011) Regulatory response to carbon starvation in Caulobacter crescentus. PLoS One 6: e18179

Carrasco B, Seco EM, Lopez-Sanz M, Alonso JC, Ayora S (2018) Bacillus subtilis RarA modulates replication restart. Nucleic Acids Res 46: 7206–7220

Chiaramello AE, Zyskind JW (1990) Coupling of DNA replication to growth rate in Escherichia coli: a possible role for guanosine tetraphosphate. J Bacteriol 172: 2013–2019

Erzberger JP, Berger JM (2006) Evolutionary relationships and structural mechanisms of AAA+ proteins. Annu Rev Biophys Biomol Struct 35: 93–114

Evinger M, Agabian N (1977) Envelope-associated nucleoid from Caulobacter crescentus stalked and swarmer cells. J Bacteriol 132: 294–301

Felletti M, Omnus DJ, Jonas K (2019) Regulation of the replication initiator DnaA in Caulobacter crescentus. Biochim Biophys Acta Gene Regul Mech 1862: 697–705

Figge RM, Divakaruni AV, Gober JW (2004) MreB, the cell shape-determining bacterial actin homologue, co-ordinates cell wall morphogenesis in Caulobacter crescentus. Mol Microbiol 51: 1321–1332

Giramma CN, DeFoer MB, Wang JD (2021) The Alarmone (p)ppGpp Regulates Primer Extension by Bacterial Primase. J Mol Biol 433: 167189

Goedhart J, Luijsterburg MS (2020) VolcaNoseR is a web app for creating, exploring, labeling and sharing volcano plots. Sci Rep 10: 20560

Gonzalez D, Collier J (2014) Effects of (p)ppGpp on the progression of the cell cycle of Caulobacter crescentus. J Bacteriol 196: 2514–2525

Gorbatyuk B, Marczynski GT (2001) Physiological consequences of blocked Caulobacter crescentus dnaA expression, an essential DNA replication gene. Mol Microbiol 40: 485–497

Gorbatyuk B, Marczynski GT (2005) Regulated degradation of chromosome replication proteins DnaA and CtrA in Caulobacter crescentus. Mol Microbiol 55: 1233–1245

Gropp M, Strausz Y, Gross M, Glaser G (2001) Regulation of Escherichia coli RelA requires oligomerization of the C-terminal domain. J Bacteriol 183: 570–579

Harinarayanan R, Murphy H, Cashel M (2008) Synthetic growth phenotypes of Escherichia coli lacking ppGpp and transketolase A (tktA) are due to ppGpp-mediated transcriptional regulation of tktB. Mol Microbiol 69: 882–894

Hauryliuk V, Atkinson GC, Murakami KS, Tenson T, Gerdes K (2015) Recent functional insights into the role of (p)ppGpp in bacterial physiology. Nat Rev Microbiol 13: 298–309

Hishida T, Ohno T, Iwasaki H, Shinagawa H (2002) Saccharomyces cerevisiae MGS1 is essential in strains deficient in the RAD6-dependent DNA damage tolerance pathway. EMBO J 21: 2019–2029

Hishida T, Ohya T, Kubota Y, Kamada Y, Shinagawa H (2006) Functional and physical interaction of yeast Mgs1 with PCNA: impact on RAD6-dependent DNA damage tolerance. Mol Cell Biol 26: 5509–5517

Irving SE, Choudhury NR, Corrigan RM (2021) The stringent response and physiological roles of (pp)pGpp in bacteria. Nat Rev Microbiol 19: 256–271

Iyer LM, Leipe DD, Koonin EV, Aravind L (2004) Evolutionary history and higher order classification of AAA+ ATPases. J Struct Biol 146: 11–31

Janion C (2008) Inducible SOS response system of DNA repair and mutagenesis in Escherichia coli. Int J Biol Sci 4: 338–344

Johnson A, O’Donnell M (2005) Cellular DNA replicases: components and dynamics at the replication fork. Annu Rev Biochem 74: 283–315

Kamat SS, Camara K, Parsons WH, Chen DH, Dix MM, Bird TD, Howell AR, Cravatt BF (2015) Immunomodulatory lysophosphatidylserines are regulated by ABHD16A and ABHD12 interplay. Nat Chem Biol 11: 164–171

Karimova G, Pidoux J, Ullmann A, Ladant D (1998) A bacterial two-hybrid system based on a reconstituted signal transduction pathway. Proc Natl Acad Sci U S A 95: 5752–5756

Kirkpatrick CL, Viollier PH (2012) Decoding Caulobacter development. FEMS Microbiol Rev 36: 193–205

Lesley JA, Shapiro L (2008) SpoT regulates DnaA stability and initiation of DNA replication in carbon-starved Caulobacter crescentus. J Bacteriol 190: 6867–6880

Leslie DJ, Heinen C, Schramm FD, Thuring M, Aakre CD, Murray SM, Laub MT, Jonas K (2015) Nutritional Control of DNA Replication Initiation through the Proteolysis and Regulated Translation of DnaA. PLoS Genet 11: e1005342

Lewis JS, Jergic S, Dixon NE (2016) The E. coli DNA Replication Fork. Enzymes 39: 31–88

Maffeo C, Chou HY, Aksimentiev A (2019) Molecular Mechanisms of DNA Replication and Repair Machinery: Insights from Microscopic Simulations. Adv Theory Simul 2

Maslowska KH, Makiela-Dzbenska K, Fijalkowska IJ (2019) The SOS system: A complex and tightly regulated response to DNA damage. Environ Mol Mutagen 60: 368–384

Miller JH (1992) A short course in bacterial genetics: a laboratory manual and handbook for Escherichia coli and related bacteria. Cold Spring Harbor Laboratory Press, Plainview, N.Y.

Modell JW, Hopkins AC, Laub MT (2011) A DNA damage checkpoint in Caulobacter crescentus inhibits cell division through a direct interaction with FtsW. Genes Dev 25: 1328–1343

Narayanan S, Janakiraman B, Kumar L, Radhakrishnan SK (2015) A cell cycle-controlled redox switch regulates the topoisomerase IV activity. Genes Dev 29: 1175–1187

Narayanan S, Kumar L, Radhakrishnan SK (2018) Sensory domain of the cell cycle kinase CckA regulates the differential DNA binding of the master regulator CtrA in Caulobacter crescentus. Biochim Biophys Acta Gene Regul Mech 1861: 952–961

Page AN, George NP, Marceau AH, Cox MM, Keck JL (2011) Structure and biochemical activities of Escherichia coli MgsA. J Biol Chem 286: 12075–12085

Puchades C, Sandate CR, Lander GC (2020) The molecular principles governing the activity and functional diversity of AAA+ proteins. Nat Rev Mol Cell Biol 21: 43–58

Quon KC, Yang B, Domian IJ, Shapiro L, Marczynski GT (1998) Negative control of bacterial DNA replication by a cell cycle regulatory protein that binds at the chromosome origin. Proc Natl Acad Sci U S A 95: 120–125

Riber L, Lobner-Olesen A (2020) Inhibition of Escherichia coli chromosome replication by rifampicin treatment or during the stringent response is overcome by de novo DnaA protein synthesis. Mol Microbiol 114: 906–919

Ronneau S, Hallez R (2019) Make and break the alarmone: regulation of (p)ppGpp synthetase/hydrolase enzymes in bacteria. FEMS Microbiol Rev 43: 389–400

Ryan KR, Judd EM, Shapiro L (2002) The CtrA response regulator essential for Caulobacter crescentus cell-cycle progression requires a bipartite degradation signal for temporally controlled proteolysis. J Mol Biol 324: 443–455

Sanselicio S, Berge M, Theraulaz L, Radhakrishnan SK, Viollier PH (2015) Topological control of the Caulobacter cell cycle circuitry by a polarized single-domain PAS protein. Nature Communications 6

Saugar I, Parker JL, Zhao S, Ulrich HD (2012) The genome maintenance factor Mgs1 is targeted to sites of replication stress by ubiquitylated PCNA. Nucleic Acids Res 40: 245–257

Schindelin J, Arganda-Carreras I, Frise E, Kaynig V, Longair M, Pietzsch T, Preibisch S, Rueden C, Saalfeld S, Schmid B et al (2012) Fiji: an open-source platform for biological-image analysis. Nat Methods 9: 676–682

Schrader JM, Li GW, Childers WS, Perez AM, Weissman JS, Shapiro L, McAdams HH (2016) Dynamic translation regulation in Caulobacter cell cycle control. Proc Natl Acad Sci U S A 113: E6859–E6867

Sekimizu K, Bramhill D, Kornberg A (1987) ATP activates dnaA protein in initiating replication of plasmids bearing the origin of the E. coli chromosome. Cell 50: 259–265

Sherratt DJ, Soballe B, Barre FX, Filipe S, Lau I, Massey T, Yates J (2004) Recombination and chromosome segregation. Philos Trans R Soc Lond B Biol Sci 359: 61–69

Simonsen S, Sogaard CK, Olsen JG, Otterlei M, Kragelund BB (2024) The bacterial DNA sliding clamp, beta-clamp: structure, interactions, dynamics and drug discovery. Cell Mol Life Sci 81: 245

Sinha AK, Lobner-Olesen A, Riber L (2020) Bacterial Chromosome Replication and DNA Repair During the Stringent Response. Front Microbiol 11: 582113

Siwach M, Kumar L, Palani S, Muraleedharan S, Panis G, Fumeaux C, Mony BM, Sanyal S, Viollier PH, Radhakrishnan SK (2021) An organelle-tethering mechanism couples flagellation to cell division in bacteria. Dev Cell 56: 657–670 e654

Skerker JM, Laub MT (2004) Cell-cycle progression and the generation of asymmetry in Caulobacter crescentus. Nat Rev Microbiol 2: 325–337

Stott KV, Wood SM, Blair JA, Nguyen BT, Herrera A, Mora YG, Cuajungco MP, Murray SR (2015) (p)ppGpp modulates cell size and the initiation of DNA replication in Caulobacter crescentus in response to a block in lipid biosynthesis. Microbiology (Reading) 161: 553–564

Taylor JA, Ouimet MC, Wargachuk R, Marczynski GT (2011) The Caulobacter crescentus chromosome replication origin evolved two classes of weak DnaA binding sites. Mol Microbiol 82: 312–326

Thanbichler M, Iniesta AA, Shapiro L (2007) A comprehensive set of plasmids for vanillate- and xylose-inducible gene expression in Caulobacter crescentus. Nucleic Acids Res 35: e137

Travis BA, Schumacher MA (2022) Diverse molecular mechanisms of transcription regulation by the bacterial alarmone ppGpp. Mol Microbiol 117: 252–260

Voedts H, Anoyatis-Pele C, Langella O, Rusconi F, Hugonnet JE, Arthur M (2024) (p)ppGpp modifies RNAP function to confer beta-lactam resistance in a peptidoglycan-independent manner. Nat Microbiol 9: 647–656

Wang B, Grant RA, Laub MT (2020) ppGpp Coordinates Nucleotide and Amino-Acid Synthesis in E. coli During Starvation. Mol Cell 80: 29–42 e10

Wang JD, Sanders GM, Grossman AD (2007) Nutritional control of elongation of DNA replication by (p)ppGpp. Cell 128: 865–875

Yao NY, O’Donnell ME (2021) The DNA Replication Machine: Structure and Dynamic Function. Subcell Biochem 96: 233–258

Yoshimura A, Seki M, Enomoto T (2017) The role of WRNIP1 in genome maintenance. Cell Cycle 16: 515–521

