## Supplemental Information for "A starvation-triggered AAA+ ATPase halts chromosome replication progression by disassembling the bacterial DNA sliding clamp"

### SUPPLEMENTAL FIGURES

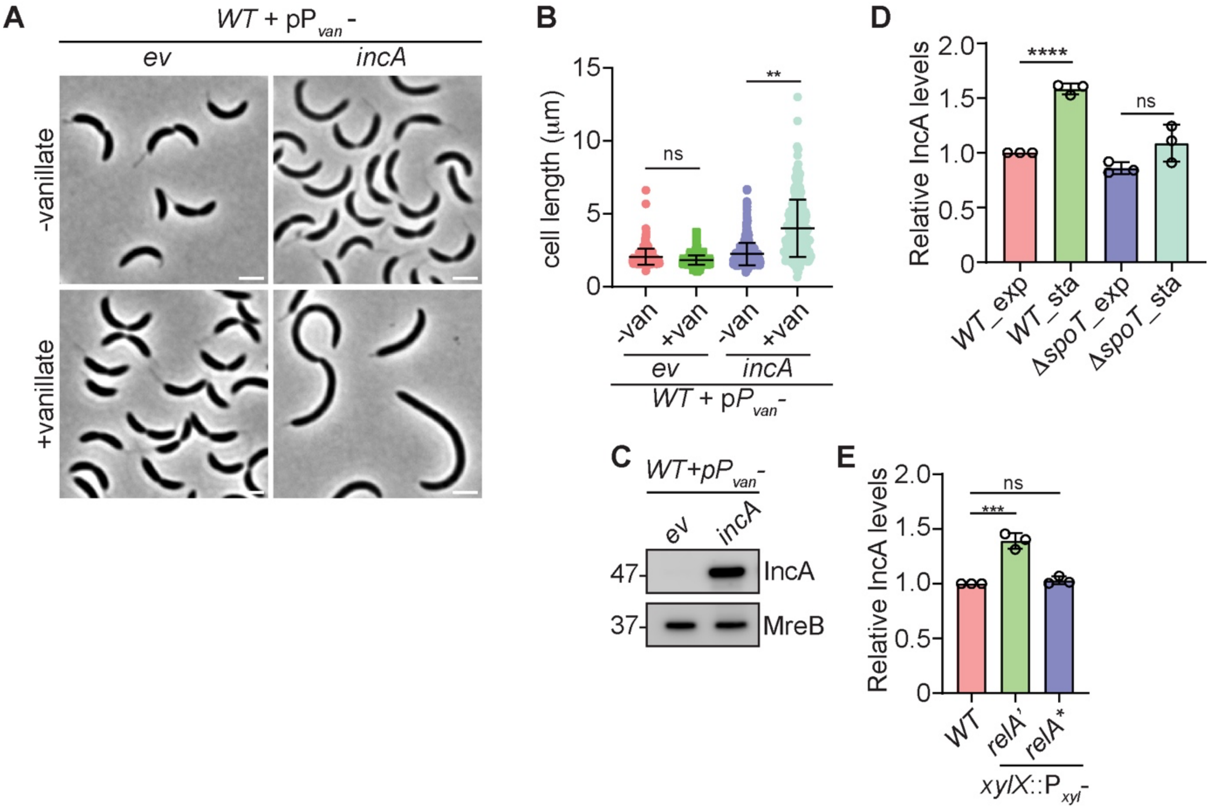


**Figure S1.** IncA production is dependent on (p)ppGpp. **(A)** Phase contrast micrographs of wild-type (*WT*) *Caulobacter* cells harbouring the high copy vector pBVMCS-4 or ectopically expressing *incA* from the vanillate inducible promoter ($P_{van}$) on pBVMCS-4. The cells were gown in the presence of 0.5mM vanillate for 5h to induce the production of *incA*. **(B)** Cell size distribution of cells from (A). Mean cell size of at least 150 cells from each biological replicate were used for the statistical analyses. **(C)** Immunoblots showing the protein levels of IncA and MreB in cells from (A). IncA protein level, relative to *WT*, in **(D)** *WT* and *spoT* null mutant (Δ*spoT*) at exponential (exp) and stationary (sta) phase and **(E)** Exponential stage *WT* cells expressing *relA'* or *relA** from the chromosomal *xylX* locus (*xylX*::$P_{xyl}$). The error bars in (B, D and E) represent the mean ± SD from at least three independent biological replicates. Statistical analyses were done using unpaired two-tailed t test; ****$p < 0.0001$, ***$p < 0.001$, **$p < 0.01$; ns, not significant. Scale bar: 2μm.

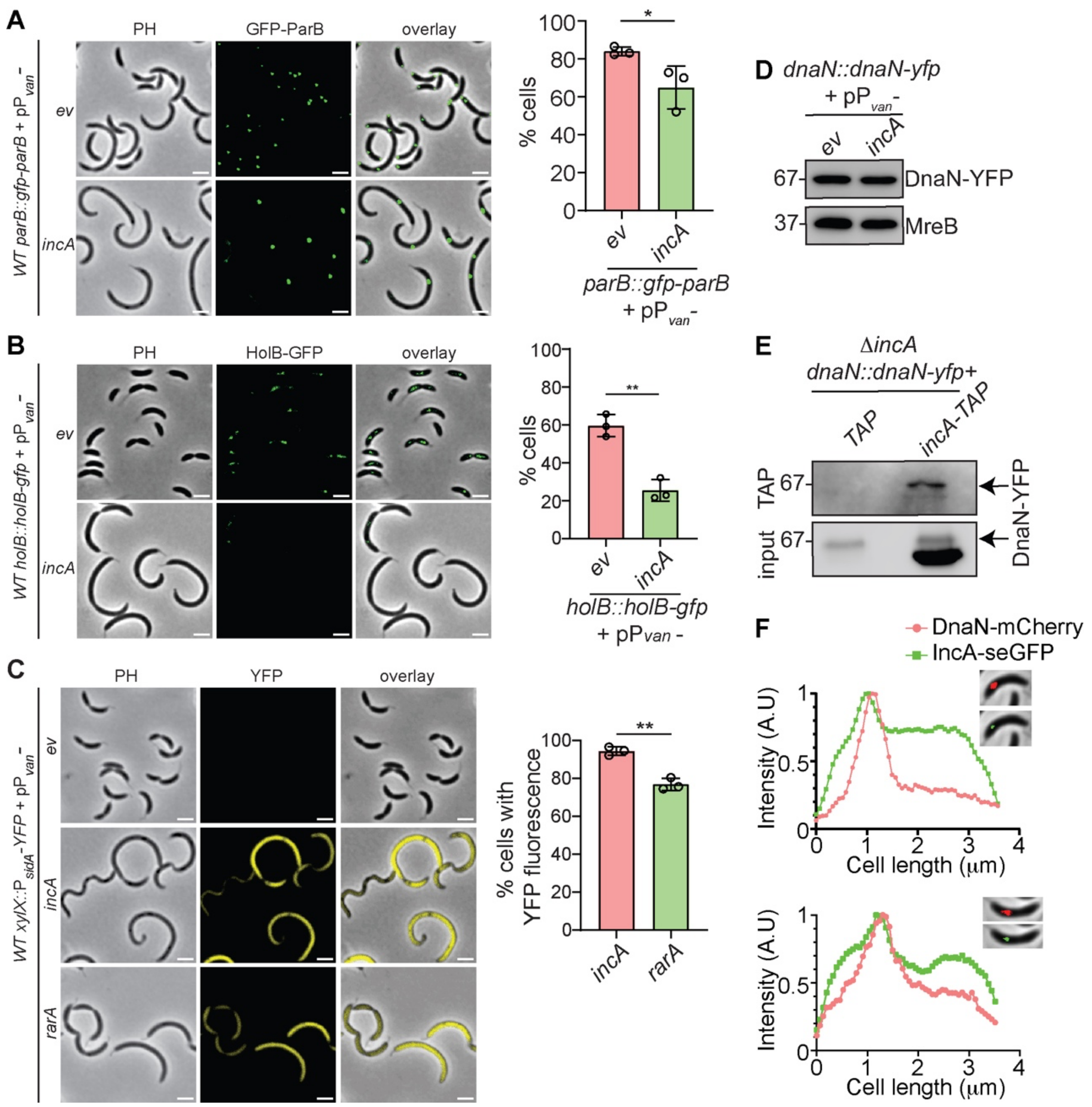


**Figure S2**. IncA inhibits chromosome replication by interacting with DnaN. Phase contrast and fluorescence micrographs of wild-type (*WT*)cells expressing **(A)** GFP-ParB from the native *parB* locus (*parB*::*gfp-parB*), **(B)** HolB-GFP from the native *holB* locus (*holB*::*holB-gfp*) and **(C)** YFP from the SOS-inducible *sidA* promoter ($P_{sidA}$) at the xylose locus (xylX::$P_{sidA}$-YFP) in cells overexpressing *incA* or *rarA* from the vanillate-inducible promoter on the high copy vector pBVMCS-4 ($pP_{van}$). The cells were induced with 0.5mM vanillate for 5 hours. Percentage of cells displaying two ParB foci or proper localization of HolB-GFP or expressing YFP are represented on the right of the respective micrographs. The error bars represent mean ± SD from at least three independent biological replicates. At least one hundred cells from each biological replicate was used for the analyses. Statistical analyses were done using unpaired two-tailed t test. Immunoblots denoting **(D)** DnaN-YFP and MreB protein levels in *WT Caulobacter* cells harbouring pBVMCS-4 or overexpressing IncA from

$P_{van}$ on pBVMCS-4 and **(E)** DnaN-YFP in tandem affinity purification (TAP) samples from Δ*incA* cells expressing *dnaN*-*yfp* from the native *dnaN* locus (*dnaN*::*dnaN*-*yfp*) and expressing TAP-tagged IncA (IncA-TAP) or the TAP-tag alone. **(F)** Fluorescence profiles of co-localised DnaN-mCherry and IncA-sEGFP in *WT* cells. ****$p < 0.0001$, **$p < 0.01$; *$p < 0.05$; ns, not significant. Scale bar: 2μm.

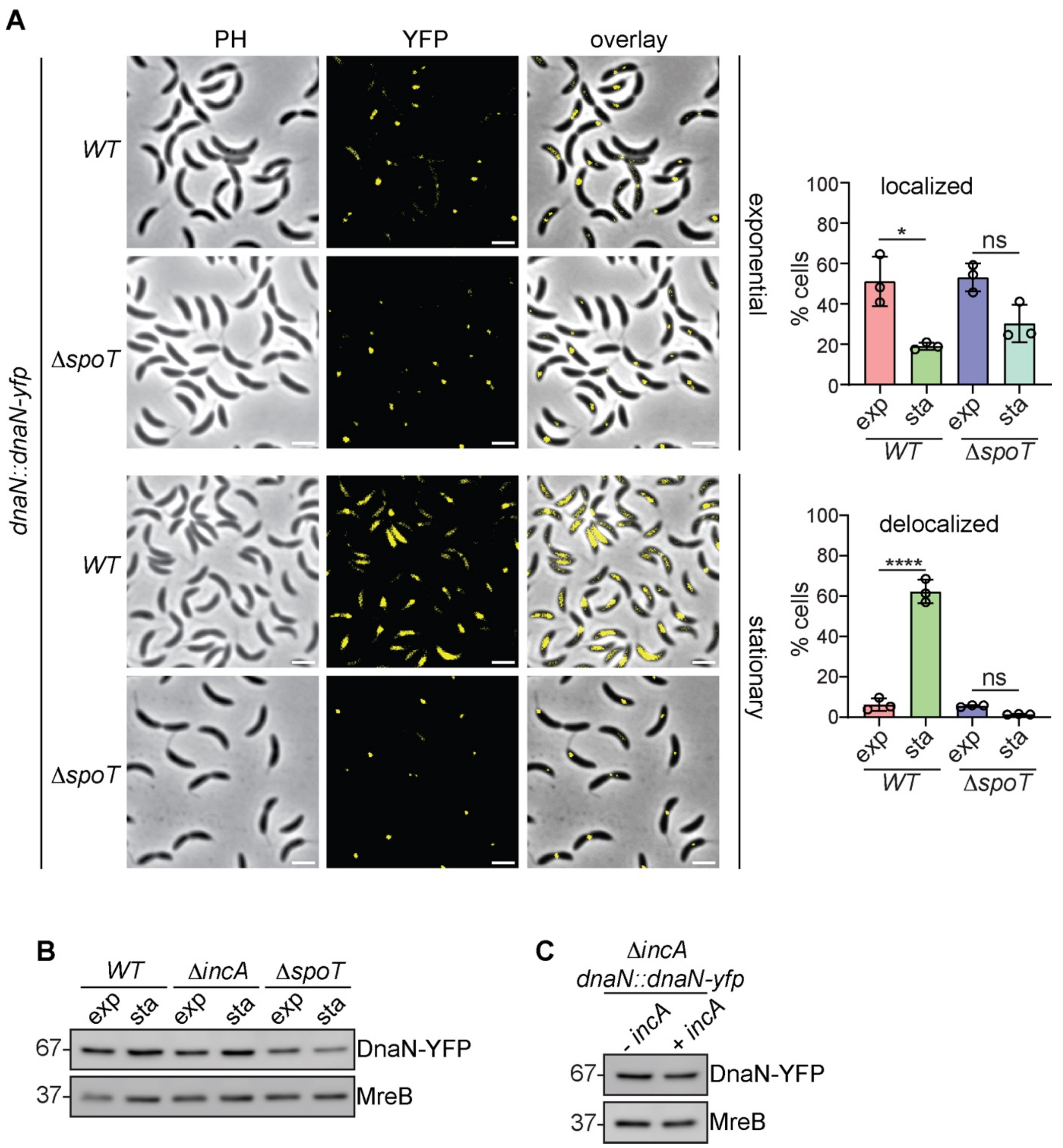


**Figure S3.** IncA induces DnaN delocalization in (p)ppGpp-abundant cells. **(A)** Phase contrast and fluorescence micrographs of stationary and exponential phase *WT* and *spoT* null mutant (Δ*spoT*) cells expressing DnaN-YFP from the native *dnaN* locus (*dnaN*::*dnaN-yfp*). Exponential phase cells were collected at $OD_{600}$ of 0.6 and stationary phase cells were collected six hours after the exponential phase. Quantification (%) of cells having localized or delocalized DnaN-YFP are represented on the right side of the micrographs. **(B-C)** Immunoblots showing the protein levels of DnaN-YFP and MreB (loading control) in cells used in Figures 3A, 3B, and S3A. The error bars represent mean ± SD from three independent biological replicates. At least one hundred cells from each biological replicate was used for the statistical analyses done using a two-way ANOVA with Holm-Sidak's multiple comparisons test; ****p < 0.001, *p < 0.05; ns, not significant. Scale bar: 2μm.

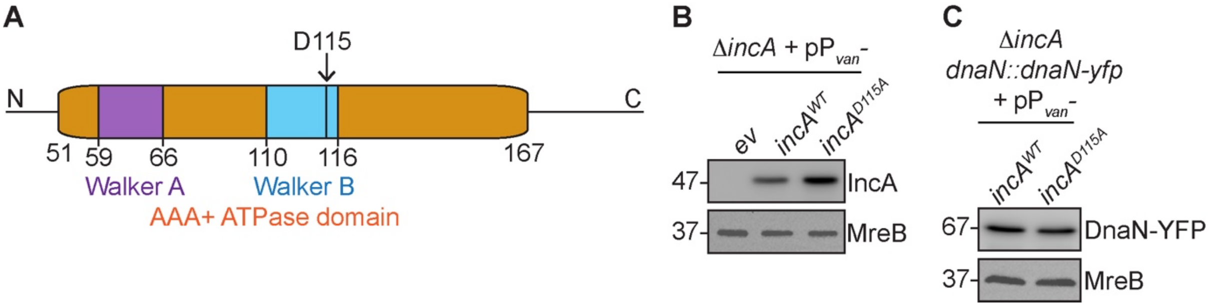


**Figure S4.** The ATPase activity of IncA is required to inhibit replication. **(A)** The domain architecture of IncA protein. The walker A (purple) and walker B (blue) motifs in the AAA+ ATPase domain (brown) are denoted. The ATP binding aspartate (D115) in the Walker B motif is also denoted. Mutation in D115 eliminates ATPase activity. Immunoblots denoting protein levels of **(B)** IncA and MreB (loading control) in cells used in Fig. 4C-E and **(C)** DnaN-YFP and MreB in cells used in Fig. 4 A. The samples for immunoblots were collected after the cells were induced with 0.5mM vanillate for 5 hours.

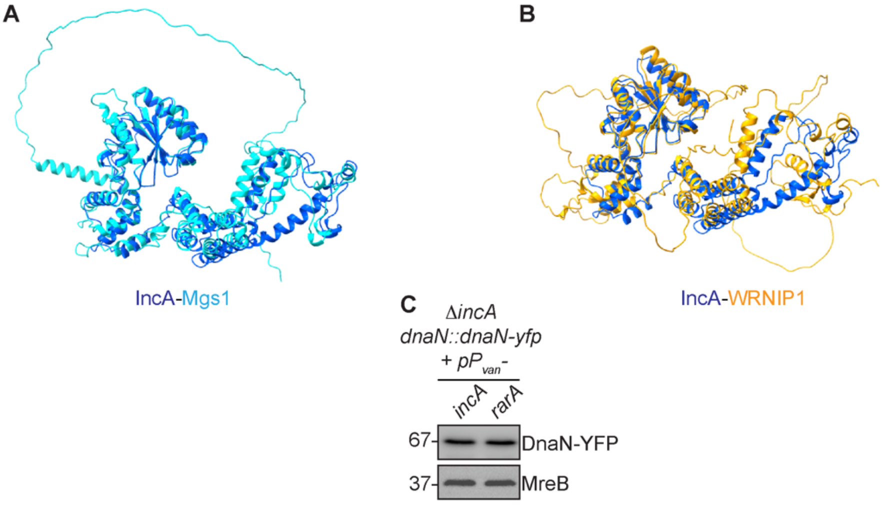


**Figure S5.** Paralogs of IncA. **(A, B)** Structural alignment of the *Caulobacter* IncA (Purple) with (A) Yeast Mgs1 (Blue) and (B) Human WRNIP1 (Orange). The monomeric protein structures were generated using AlphaFold and docked using UCSF ChimeraX. **(C)** Immunoblot showing the protein levels of DnaN-YFP and MreB (loading control) in cells used in Fig. 5A.

### SUPPLEMENTAL DATASET LEGEND

**Supplemental Dataset 1.** Quantitative mass spectrometry analyses dataset

### SUPPLEMENTAL VIDEO LEGENDS

**Supplemental Video 1.** Fluorescence live-cell imaging of DnaN-YFP in *WT Caulobacter* cells harbouring the empty vector pBVMCS-4 (*ev*). Time-lapse series were acquired at a 5 min interval. Cells were imaged at room temperature on 1% agarose in PYE supplemented with 0.5mM vanillate. The video was rendered at 4 fps in the compressed avi format for visualization purposes. Scale bar: 2µm

**Supplemental Video 2.** Fluorescence live-cell imaging of DnaN-YFP in *WT Caulobacter* cells overexpressing *incA* from a vanillate-inducible promoter on pBVMCS-4 (p$P_{van}$-*incA*). Time-lapse series were acquired at a 5 min interval. Cells were imaged at room temperature on 1% agarose in PYE supplemented with 0.5mM vanillate. The video was rendered at 4 fps in the compressed avi format for visualization purposes. Scale bar: 2µm

### SUPPLEMENTAL MATERIALS AND METHODS

#### Promoter activity assay

*Caulobacter* strains harbouring the promoter of *incA* fused to the lacZ reporter gene ($P_{incA}$-*lacZ*) were grown at 29 °C till it reached $OD_{660}$ of 0.2-0.4 ($Ab_{660}$). From this, 50 µl of cells were lysed using 10 µl of chloroform to which 750 µl of Z-buffer (60 mM $Na_2HPO_4$, 40 mM $NaH_2PO_4.H_2O$, 10 mM KCl, 1 mM $MgSO_4.7H_2O$, pH-7) was added, followed by vigorous vortexing. To this lysed mixture 200 µL of Ortho Nitro Phenyl-β-D Galactoside (4 mg $ml^{-1}$ ONPG in 0.1 M potassium phosphate buffer pH-7) was added. The reaction mixture was then incubated at 30 °C till a yellow colour developed and the reaction was stopped with the addition of 500 µL of 1 M $Na_2CO_3$. Finally, the absorbance of the reaction mixture was measured at 420 nm using Z-buffer as blank. The lacZ activity (Miller units) was calculated using the following equation, U = ($Ab_{420}$*1000) / ($Ab_{660}$*t*V), where t is time taken to develop the yellow colour in min, V is the volume of cells in ml.

**Structural alignments**

For the protein structural alignment, AlphaFold (Jumper *et al*, 2021; Varadi *et al*, 2024) predicted structures of *C. crescentus* IncA (AlphaFold id: AF-A0A0H3C719-F1), *E. coli* RarA (AlphaFold id: AF-P0AAZ4-F1), *S. cerevisiae* Mgs1 (AlphaFold id: AF-P40151-F1) and Human WRNIP1 (AlphaFold id: AF-Q96S55-F1) were used to generate structural alignments using UCSF ChimeraX(Goddard *et al*, 2018; Pettersen *et al*, 2021).

**SUPPLEMENTAL TABLES**

**Table S1: *Caulobacter crescentus* strains used in this study**

| Identifier | Strain/ Genotype | Source |
|---|---|---|
| NA1000 | *Wild-type Caulobacter crescentus* | (Evinger & Agabian, 1977) |
| SY466 | Δ*incA* | This study |
| SY801 | *WT* + pBVMCS-4 | This study |
| SY803 | *WT* + pBVMCS-4-P$_{van}$-*incA* | This study |
| NABC124 | *WT dnaN::dnaN-yfp* | (Joseph *et al*, 2021) |
| SY234 | *WT dnaN::dnaN-yfp* + pBVMCS-4 | This study |
| SY235 | *WT dnaN::dnaN-yfp* + pBVMCS-4-P$_{van}$-*incA* | This study |
| SY558 | Δ*spoT* | (Stott *et al*, 2015) |
| JC820 | *WT* P$_{xyl}$*_relA'* | (Gonzalez & Collier, 2014) |
| JC1198 | *WT* P$_{xyl}$*_relA** | (Gonzalez & Collier, 2014) |
| SY544 | *WT dnaN::dnaN-yfp* | This study |
| SY547 | Δ*incA dnaN::dnaN-yfp* | This study |
| SY655 | Δ*spoT dnaN::dnaN-yfp* | This study |
| SY692 | Δ*incA dnaN::dnaN-yfp* + pRKlac290 | This study |
| SY694 | Δ*incA dnaN::dnaN-yfp* + RKlac290-P$_{incA}$-*incA* | This study |
| SY649 | *WT* + pRKlac290-P$_{incA}$-*lacZ* | This study |
| SY651 | Δ*spoT* + pRKlac290-P$_{incA}$-*lacZ* | This study |
| SY700 | *WT* P$_{xyl}$-*relA'* + pRKlac290-P$_{incA}$-*lacZ* | This study |
| SY702 | *WT* P$_{xyl}$-*relA** + pRKlac290-P$_{incA}$-*lacZ* | This study |
| SY125 | *WT dnaN::dnaN-mCherry* | This study |
| SY717 | *WT dnaN::dnaN-mCherry* + pBVMCS-4-P$_{van}$-*incA-segfp* | This study |
| AKS393 | Δ*incA dnaN::dnaN-YFP* + pBXMCS-4-TAP | This study |
| AKS353 | Δ*incA dnaN::dnaN-YFP* + pBVMCS-4-*incA*-TAP | This study |
| MT174 | *WT gfp::gfp-ParB* | (Thanbichler & Shapiro, 2006) |
| SY981 | *WT gfp::gfp-ParB* + pBVMCS-4 | This study |
| SY983 | *WT gfp::gfp-ParB* + pBVMCS-4-P$_{van}$-*incA* | This study |
| SY809 | Δ*incA* + pBVMCS-4 | This study |
| SY811 | Δ*incA* + pBVMCS-4-P$_{van}$-*incA* | This study |
| SY815 | Δ*incA* + pBVMCS-4-P$_{van}$-*incA$^{D115A}$* | This study |
| SY996 | Δ*incA* + pBVMCS-4-P$_{van}$-*rarA* | This study |
| SY991 | Δ*incA dnaN::dnaN-yfp* + pBVMCS-4-P$_{van}$-*incA* | This study |
| SY835 | Δ*incA dnaN::dnaN-yfp* + pBVMCS-4-P$_{van}$-*incA$^{D115A}$* | This study |
| SY837 | Δ*incA dnaN::dnaN-yfp* + pBVMCS-4-P$_{van}$-*rarA* | This study |
| *NABC581* | *WT xylX::*P$_{sidA}$-*yfp* | (Joseph *et al*, 2022) |
| SY959 | *WT xylX::*P$_{sidA}$-*yfp* + pBVMCS-4 | This study |
| SY961 | *WT xylX::*P$_{sidA}$-*yfp* + pBVMCS-4-P$_{van}$-*incA* | This study |
| SY963 | *WT xylX::*P$_{sidA}$-*yfp* + pBVMCS-4-P$_{van}$-*rarA* | This study |
|  | *WT holB:: holB-gfp* | (Jensen *et al*, 2001) |
| SY912 | *WT holB:: holB-gfp* + pBVMCS-4 | This study |
| SY914 | *WT holB:: holB-gfp* + pBVMCS-4-P$_{van}$-*incA* | This study |

**Table S2: *Escherichia coli* strains used in this study**

| Identifier | Strain/ Genotype | Source |
|---|---|---|
| EC100D | *F- mcrA Δ(mrr-hsdRMS-mcrBC) Φ80dlacZΔM15 ΔlacX74 recA1 endA1 araD139 Δ(ara, leu)7697 galU galK λ- rpsL nupG* | Lucigen |
| BTH101 | F- , cya-99, araD139, galE15, galK16, rpsL1 (Str r ), hsdR2, mcrA1, mcrB1 | Euromedex |
| SY690 | BTH101 + pKT25-*zip* + pUT18C-*zip* | This study |
| SY698 | BTH101 + pKT25-*incA* + pUT18 | This study |
| SY671 | BTH101 + pKT25-*incA* + pUT18-*dnaN* | This study |
| SY672 | BTH101 + pKT25-*incA* + pUT18-*holB* | This study |
| SY673 | BTH101 + pKT25-*incA* + pUT18-*ssb* | This study |
| SY864 | BTH101 + pKT25-*rarA* + pUT18-*dnaN* | This study |
| SY908 | BTH101 + pKT25-$incA^{D115A}$ + pUT18-*dnaN* | This study |

**Table S3: Plasmids used in this study**

| **Plasmids** | **Description** | **Source** |
|---|---|---|
| pSY27 | pBVMCS-4; $Gent^R$ | (Thanbichler *et al*, 2007) |
| pSY01 | pBVMCS-4-$P_{van}$-*incA* | This study |
| pSY02 | pBVMCS-4-$P_{van}$-*incA-segfp* | This study |
| pSY191 | pRKlac290 low copy vector containing promoter-less lacZ gene; $Tet^R$ | (Gober & Shapiro, 1992) |
| pSY276 | pRKlac290-$P_{incA}$-*incA* | This study |
| pSY135 | pRKlac290-$P_{incA}$-*lacZ* | This study |
| pAKS172 | pBXMCS-4-$P_{xyl}$-*TAP* | Lab stock |
| pSKR344 | pBVMCS-4-$P_{van}$-*incA-TAP* | This study |
| pSY725 | pBVMCS-4-$P_{van}$-$incA^{D115A}$ | This study |
| pSY729 | pBVMCS-4-$P_{van}$-*rarA* | This study |
| pSY889 | pBAD24; $Amp^R$ | (Guzman *et al*, 1995) |
| pSY1172 | pBAD24-$P_{ara}$-*incA* | This study |
| pSY1173 | pBAD24-$P_{ara}$-*rarA* | This study |
| pNPTS138 | *sacB* containing suicide vector used for gene replacement; $Kan^R$ | M.R.K Alley, unpublished |
| pSKR331 | pNPTS138 derivative carrying the in-frame deletion of *incA;* $Kan^R$ | This study |
| pKT25 | Low copy plasmid with T25 C-terminus adenylate cyclase fragment; $Kan^R$ | Euromedex |
| pUT18 | High copy plasmid with T18 N-terminus adenylate cyclase fragment; $Amp^R$ | Euromedex |
| pKT25-zip | pKT25 containing the leucine zipper of GCN4 fused to the T25 fragment | Euromedex |
| pUT18C-zip | pUT18C containing the leucine zipper of GCN4 fused to the T18 fragment | Euromedex |
| pSY1174 | pKT25-*incA* | This study |
| pSY1175 | pKT25-$incA^{D115A}$ | This study |
| pSY1179 | pKT25-*rarA* | This study |
| pSY1176 | pUT18-*dnaN* | This study |
| pSY1177 | pUT18-*holB* | This study |
| pSY1178 | pUT18-*ssb* | This study |

### SUPPLEMENTAL REFERENCE

Evinger M, Agabian N (1977) Envelope-associated nucleoid from Caulobacter crescentus stalked and swarmer cells. *J Bacteriol* 132: 294-301
Gober JW, Shapiro L (1992) A developmentally regulated Caulobacter flagellar promoter is activated by 3' enhancer and IHF binding elements. *Mol Biol Cell* 3: 913-926
Goddard TD, Huang CC, Meng EC, Pettersen EF, Couch GS, Morris JH, Ferrin TE (2018) UCSF ChimeraX: Meeting modern challenges in visualization and analysis. *Protein Sci* 27: 14-25
Gonzalez D, Collier J (2014) Effects of (p)ppGpp on the progression of the cell cycle of Caulobacter crescentus. *J Bacteriol* 196: 2514-2525
Guzman LM, Belin D, Carson MJ, Beckwith J (1995) Tight regulation, modulation, and high-level expression by vectors containing the arabinose PBAD promoter. *J Bacteriol* 177: 4121-4130
Jensen RB, Wang SC, Shapiro L (2001) A moving DNA replication factory in Caulobacter crescentus. *Embo J* 20: 4952-4963
Joseph AM, Daw S, Sadhir I, Badrinarayanan A (2021) Coordination between nucleotide excision repair and specialized polymerase DnaE2 action enables DNA damage survival in non-replicating bacteria. *Elife* 10
Joseph AM, Nahar K, Daw S, Hasan MM, Lo R, Le TBK, Rahman KM, Badrinarayanan A (2022) Mechanistic insight into the repair of C8-linked pyrrolobenzodiazepine monomer-mediated DNA damage. *RSC Med Chem* 13: 1621-1633
Jumper J, Evans R, Pritzel A, Green T, Figurnov M, Ronneberger O, Tunyasuvunakool K, Bates R, Zidek A, Potapenko A *et al* (2021) Highly accurate protein structure prediction with AlphaFold. *Nature* 596: 583-589
Pettersen EF, Goddard TD, Huang CC, Meng EC, Couch GS, Croll TI, Morris JH, Ferrin TE (2021) UCSF ChimeraX: Structure visualization for researchers, educators, and developers. *Protein Sci* 30: 70-82
Stott KV, Wood SM, Blair JA, Nguyen BT, Herrera A, Mora YG, Cuajungco MP, Murray SR (2015) (p)ppGpp modulates cell size and the initiation of DNA replication in Caulobacter crescentus in response to a block in lipid biosynthesis. *Microbiology (Reading)* 161: 553-564
Thanbichler M, Iniesta AA, Shapiro L (2007) A comprehensive set of plasmids for vanillate- and xylose-inducible gene expression in Caulobacter crescentus. *Nucleic Acids Res* 35: e137
Thanbichler M, Shapiro L (2006) MipZ, a spatial regulator coordinating chromosome segregation with cell division in Caulobacter. *Cell* 126: 147-162
Varadi M, Bertoni D, Magana P, Paramval U, Pidruchna I, Radhakrishnan M, Tsenkov M, Nair S, Mirdita M, Yeo J *et al* (2024) AlphaFold Protein Structure Database in 2024: providing structure coverage for over 214 million protein sequences. *Nucleic Acids Res* 52: D368-D375
